# How persistent infection overcomes peripheral tolerance mechanisms to cause T cell-mediated autoimmune disease

**DOI:** 10.1101/2023.09.13.557414

**Authors:** Rose Yin, Samuel Melton, Eric Huseby, Mehran Kardar, Arup K. Chakraborty

## Abstract

T cells help orchestrate immune responses to pathogens, and their aberrant regulation can trigger autoimmunity. Recent studies highlight that a threshold number of T cells (a quorum) must be activated in a tissue to mount a functional immune response. These collective effects allow the T cell repertoire to respond to pathogens while suppressing autoimmunity due to circulating autoreactive T cells. Our computational studies show that increasing numbers of pathogenic peptides targeted by T cells during persistent or severe viral infections increase the probability of activating T cells that are weakly reactive to self-antigens (molecular mimicry). These T cells are easily re-activated by the self-antigens and contribute to exceeding the quorum threshold required to mount autoimmune responses. Rare peptides that activate many T cells are sampled more readily during severe/persistent infections than in acute infections, which amplifies these effects. Experiments in mice to test predictions from these mechanistic insights are suggested.

## Introduction

The key cell types that enable the adaptive immune system to mount pathogen-specific responses to a diverse and evolving world of microbes are T and B lymphocytes (T cells and B cells). Humans have billions of T cells and B cells, each of which expresses a T cell receptor (TCR) or B cell receptor (BCR) on its surface. T and B cell repertoires are characterized by an enormous diversity of TCRs and BCRs generated by VDJ recombination [1, 2, 3]. Upon infection with a pathogen, some receptors from this pool are likely to bind sufficiently strongly to molecular components of a specific pathogen resulting in T or B cell activation, which can potentially result in an adaptive immune response. For example, T cell receptors (TCRs) bind to peptides (p) derived from a pathogenic protein bound to protein products of the major histocompatibility (MHC) genes. T cell activation is triggered if the TCR-pMHC bond has a sufficiently long half-life [4, 5, 6]. Different TCRs tend to bind to different pMHC molecules with sufficiently long half-lives, thus enabling the T cell repertoire to respond specifically to diverse pathogens. At the same time, the T cell repertoire is largely tolerant to pMHC molecules where the peptide is derived from host proteins. Such self-pMHC molecules are expressed ubiquitously on host cells. Tolerance to self is due to processes that occur during T cell development in the thymus and mechanisms that suppress autoimmune responses in peripheral tissues [7, 8, 9, 10].

Cells in the thymus (especially those in the medulla) express the AIRE gene which enables promiscuous gene expression and results in these cells displaying pMHC molecules with peptides derived from diverse regions of the host proteome [11]. During development, immature T cells (thymocytes) interact with these self-pMHC molecules. To successfully develop into a mature T cell, a thymocyte must bind to at least one self-pMHC molecule it encounters in the thymus with a binding free energy (or half-life) exceeding a threshold in order to receive a survival signal (positive selection) [12]. However, if a thymocyte’s TCR binds to any encountered host pMHC molecule with a binding free energy exceeding a higher threshold, it is deleted (negative selection). In this way, positive selection aims to ensure that the mature T cell repertoire expresses TCRs with the ability to bind to pMHC complexes, while negative selection aims to delete self-reactive T cells [13, 14, 15, 16]. Theoretical and experimental studies have shown that negative selection also plays an important role in mediating the peptide-specificity of TCRs; i.e., most point mutations to a TCR’s cognate peptide abrogate recognition [17, 18, 19, 20, 21, 22]. Some of these theoretical and experimental studies also showed that the peptide contact residues on TCRs of mature T cells that undergo normal negative selection are statistically enriched in amino acids that are moderately hydrophobic. We note also that thymocytes that successfully mature but express TCRs that bind more strongly to self-pMHC molecules are more likely to differentiate into regulatory T cells (Tregs) that can suppress responses from conventional mature T cells in tissues [22].

On average, a randomly picked conventional T cell from the mature repertoire has a lower probability of being activated by a randomly picked self-pMHC molecule compared to a pathogen-derived pMHC molecule. This is because the activation threshold for mature T cells is slightly higher, but similar, to that for negative selection of thymocytes [23], and every mature T cell interacted with some fraction of the self-pMHC molecules displayed in the thymus and was not deleted by negative selection. However, this difference in probabilities of activation by self and pathogen-derived pMHC molecules is likely to be small in many cases. Furthermore, a given thymocyte does not encounter every self-peptide from its host’s proteome during thymic selection. If a mature T cell encounters a self-pMHC molecule that it did not encounter in the thymus, the difference in the probability that it will be activated by this self-pMHC molecule compared to a randomly picked pathogen-derived pMHC molecule is likely to be non-existent. These arguments suggest that imperfect thymic selection and stochastic effects associated with T cell activation should prevent robust discrimination between self and foreign antigens at the level of an individual T cell. Indeed, the mature T cell repertoire is known to include T cells that can be activated by some host-derived peptides [24, 25, 26, 27, 9, 28]. Full blown autoimmune responses to these self-peptides is thought to be suppressed by peripheral tolerance mechanisms, which include the action of Tregs [29, 30] and more recent reports of CD8^+^ suppressor T cells [8]. But, differences in the probabilities of activation of a T cell by self and pathogen-derived pMHC molecules can be small to non-existent. So, how do peripheral tolerance mechanisms suppress self-activated T cells and autoimmunity while not suppressing the ability of the repertoire to respond effectively to pathogens?

In order to mount a functional immune response, T cells must not only be activated but also proliferate and differentiate to acquire effector functions. Proliferation and differentiation require cytokines [31, 32]. For example, IL2 is necessary to promote proliferation [33, 34]. Tregs can compete with activated cells for these resources necessary for growth and differentiation to inhibit activated T cells from mounting a functional immune response [35, 36, 10]. A theoretical analysis [37] posited that if a sufficient number of T cells were activated in the same tissue, they could produce enough of the factors necessary for proliferation and differentiation and share these resources with each other to overcome the competition from suppressive regulatory mechanisms such as those mediated by Tregs. This concept provides a mechanism for how activated autoreactive T cells are inhibited from mounting a functional immune response by peripheral suppressive mechanisms while T cell activation by pathogenic pMHC molecules can overcome this effect. The difference in probability of a T cell being activated by self and pathogen-derived pMHC is amplified by the requirement that a sufficient number of T cells must be activated in the same tissue for an effective immune response to ensue. To illustrate how this collective effect amplifies this difference, as a contrived example let the average probability of a T cell being activated by a pathogen-derived and host-derived pMHC molecule be 0.5 and 0.4, respectively (1.25-fold difference); the difference is due to thymic selection. If the number of T cells that need to be activated for an effective response is 10, the probability of a functional immune response to a pathogenic or self-pMHC molecule is 0.5^10^ and 0.4^10^, respectively. So, a functional response to a pathogenic pMHC molecule is over 9 (1.25^10^) times higher than for a host-derived pMHC molecule. Due to stochastic effects, a 1.25-fold difference in the probability of T cell activation by self and pathogenic pMHC molecules is unlikely to robustly differentiate between self and non-self, but a 9-fold difference can be sufficient. Thus, collective effects can provide a robust mechanism enabling the T cell repertoire to mount effective responses against pathogens while suppressing T cells activated by self in spite of imperfect thymic selection and stochastic effects. This mechanism proposed by Butler et. al. can be considered to be akin to quorum sensing by bacteria [37, 38]. The quorum number of activated T cells required for a functional response has to be such that, with high probability, it is not exceeded upon interactions with host pMHC molecules, but is exceeded by pathogen-derived pMHC molecules [Fig. 1].

**Figure 1:**
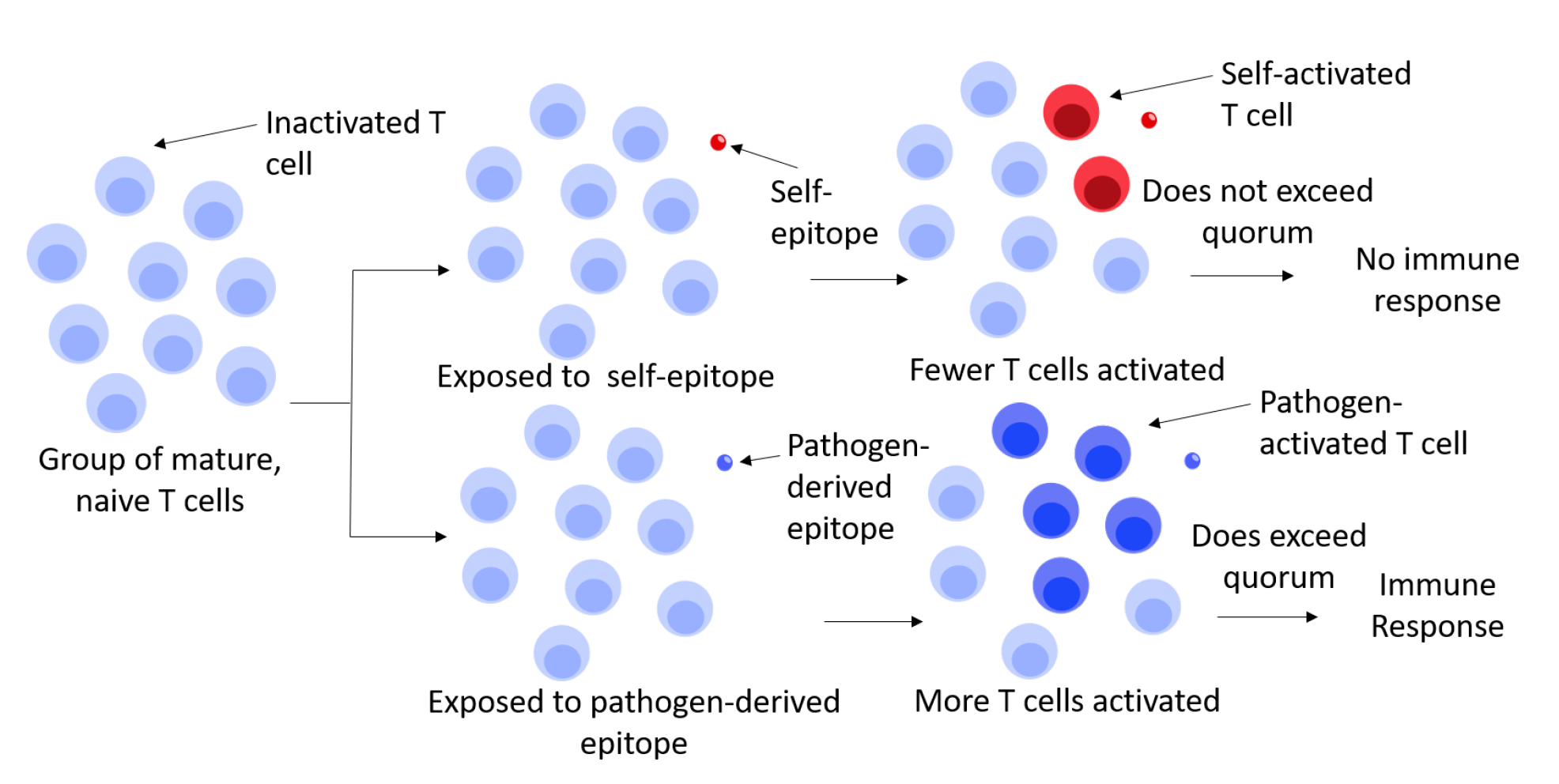
Schematic depiction of how quorum sensing by T cells can result in functional immune responses to pathogen-derived pMHC molecules and not host-derived ones in spite of the existence of autoreactive T cells in the mature repertoire. For the case depicted, the quorum number is taken to be four.

Note that the amplification of small differences in T cell activation probabilities to result in large differences in the probability of a functional response due to collective effects embodied in the quorum model is a highly non-linear effect. If the quorum number is 10, then the difference in the probability of a functional response if 5 or 6 T cells are activated is negligible, but the difference in outcome if 9 or 10 T cells are activated is very large.

Several experimental observations provide evidence for the importance of collective effects and the quorum model. Following observations of T cells forming clusters by interacting with the same APC [39] and suggestions of potential cooperativity between T cells [40, 41, 42, 43], Gerard et. al. used two-photon microscopy in mice to discover junction formation between CD8 T cells and show that these interactions increased T cell sensitivity to cytokines [44]. This study also showed that interactions between adhesion molecules LFA-1 and ICAM-1, both expressed on T cells, was important for junction formation. Junction formation mediated sharing of the cytokine, IFN-γ, and this was important for CD8 T cell differentiation and memory cell development. In a more recent study [45], related results were reported in vitro. CD8 T cell clustering around DCs expressing a stimulatory ligand was mediated by ICAM-1 binding to LFA-1 proteins on other T cells. CD80 expression also increased on activated T cells, enabling binding to CD28 on neighboring T cells. CD28 binding to CD80 results in signaling that mediates the secretion of several cytokines, including IL2 [46, 47, 48, 49]. Experiments showed that clustered T cells expand in an IL2-dependent manner and that the IL2 amount scaled with the density of clustered activated T cells.

Another elegant study combined experiments in mouse models and computation [10]. Tregs require IL2 for mediating their suppressive functions but do not secrete IL2 themselves [50, 36]. The IL2 receptor (IL2Rα) on Tregs can bind to IL2 produced by vicinal activated T cells to sequester IL2 and trigger Treg proliferation and effector functions. Wong et. al. showed that this feedback regulation of Treg’s suppressive effects could be inhibited by preventing the expression of molecules like IL2Rα in Tregs. Inhibiting Treg activity led to IL2-mediated signaling in activated T cells. The increase in signaling and IL2 production can then outcompete Tregs’ ability to bind to IL2. The range of IL2 diffusion is also thus increased, which allows other activated T cells in the vicinity to access IL2. This study also showed that Treg density controls the balance between Treg-mediated suppression and IL2 signaling by activated T cells in a non-linear way. This observation is consistent with the quorum model. Treg density should control the number of activated T cells required to outcompete the effects of Tregs (the quorum number), and the dependence should be non-linear due to the collective effects inherent in the quorum model (see above). Related observations were made in another study in mice and in vitro on the differentiation of activated CD4 T cells to memory cells [51]. Differentiation into memory T cells in mice was dependent on the precursor frequency of antigen-specific CD4 T cells. In vitro, differentiation into memory cells in microwells was density-dependent. IL2 concentrations were found to decline rapidly with distance from the secreting T cell, thus explaining the density dependence of IL2-mediated signaling that can result in differentiation. Evidence for quorum sensing is also suggested by comparing the statistical properties of the sequences of the CDR3 regions of mouse T cell repertoires at different stages of thymic development [52]. This study found that post and pre-selection repertoires could be distinguished based on the statistical properties of the ensemble of sequences rather than individual sequences, thus arguing for collective effects in self-nonself discrimination.

A T cell-mediated autoimmune condition results when the central and peripheral tolerance mechanisms described above fail. In several contexts, it has been observed that persistent pathogen infections often trigger T cell mediated autoimmunity. Examples include type I diabetes (T1D) [53, 54, 55] and multiple sclerosis (MS) [56, 57]. In the case of MS, longitudinal studies have shown that infection with Epstein-Barr Virus (EBV) is a necessary condition for MS [58] [Fig. 2A]. Recent studies suggest that T1D may be triggered by long-term infections by human enteroviruses (EV) or long COVID [59, 60, 61] [Fig. 2B]. Even though EV has typically been seen as an acute infection, it seems that EV can persistently infect human pancreatic islet cells [62] that are the targets of autoimmune responses in T1D.

**Figure 2:**
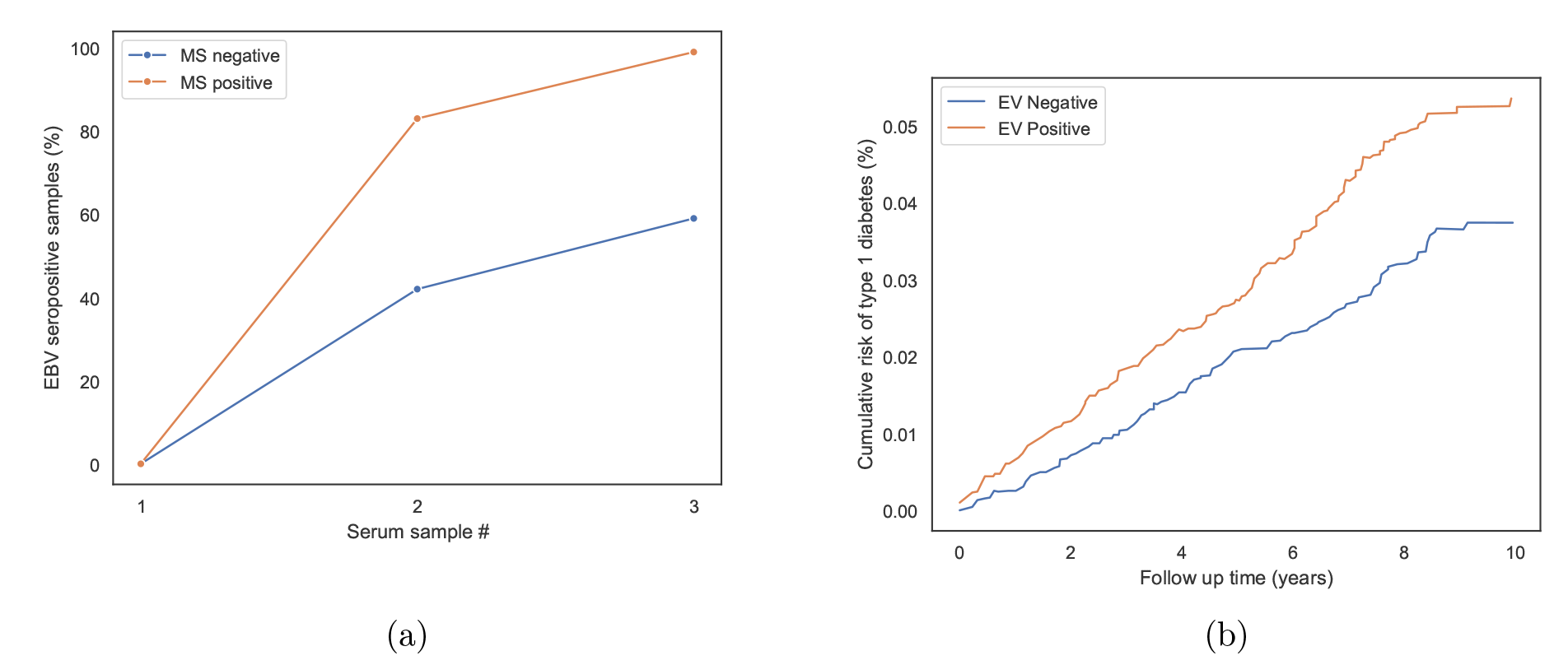
Examples of increasing autoimmunity risk given persistent infection. (a) Figure adapted from Figure 2 in Ref. [58]. This study tracked MS and EBV status of over 10 million individuals in the US military over 20 years. This figure shows the proportion of individuals with and without MS who were EBV-positive at three different times. Serum sample number was determined from the MS cohort. The 1st serum sample corresponds to the first serum sample collected from each MS patient, the 3rd sample corresponds to the one last collected before MS disease onset, and the 2^nd^ sample corresponds to one collected in between. There were two controls (MS negative) samples for each MS positive sample with the same age, sex, race/ethnicity, branch of military service, and blood sample collection dates. As seen, a significantly higher proportion of individuals who later developed MS were EBV-positive in the second and third times of serum collection compared to those who were MS-negative. (b) Figure adapted from Figure 2 in Ref. [63]. This study tracked T1D status of 570,133 children with EV and frequency-matched children without EV over 10 years. Cumulative risk of T1D increased more rapidly in children with EV than those without.

Infection-induced onset of autoimmunity has been attributed to diverse factors including specific MHC genetics, T regulatory cell dysfunction, epitope spreading, and molecular mimicry [64, 65]. The “molecular mimicry” concept suggests that T cells primed by the foreign antigen are crossreactive to similar self-peptides, thus resulting in an autoimmune response. For example, there is evidence that, compared to healthy people, MS patients exhibit higher levels of CD4+ T cells that are crossreactive to EBV peptides and those derived from myelin (a target of autoimmune responses in MS) [66, 67]. Human Leukocyte Antigen (HLA)-B27 is a human MHC allele that is associated with autoimmune diseases like ankylosing spondylitis (AS) and acute anterior uveitis (AAU). Recent studies have identified peptides derived from microbes and self-peptides presented by HLA B27 that activate T cells isolated from AS and AAU patients [67]. The “epitope spreading” mechanism proposes that an antiviral immune response leads to tissue damage and the release of otherwise sequestered self-antigens into the local region, which are then presented by nearby APCs to activate autoreactive T cells [56]. In the case of T1D, it is well known that both CD4+ and CD8+ T cells progressively destroy islet beta cells in the pancreas [68, 69, 70, 71], thus potentially generating higher levels of tissue antigens. Higher levels of inflammation in infected tissues can also result in higher levels of self-antigen presentation on antigen presenting cells [72, 73], including increased cross-priming of MHC class I responses [74].

Despite these proposed mechanisms, clear explanations for several features of the association of autoimmune diseases with persistent viral infections are not available. For example, why do the mechanisms that normally suppress autoimmunity discussed above fail for persistent or severe viral infections? Why does EBV infection only increase the probability of developing MS and not always lead to disease? In addition, individuals with the persistent viral infection do not immediately develop the corresponding autoimmune condition. This latency of pathogenesis is not sufficiently explained by the current paradigms.

Based on computational studies, we propose a mechanism for why certain autoimmune conditions are triggered by particular persistent viral infections. This mechanism is compatible with the concepts of molecular mimicry and epitope spreading, but relies critically on understanding why the collective non-linear effects embodied in the quorum model that usually suppress autoimmunity fail upon persistent or severe infections. Consistent with molecular mimicry, our computational studies suggest that specific persistent viral infections can trigger particular autoimmune conditions because T cells activated by certain peptides derived from this pathogen are weakly cross-reactive to specific self-antigens presented in a tissue. Normally, the weak reactivity to these self-antigens does not activate these T cells and so they cannot contribute to exceeding the quorum threshold for mounting functional immune responses to these antigens. Our experimental data show that the activation threshold is lower for previously activated T cells [Fig. 5]. Therefore, once activated by the pathogen-derived peptide, the activation threshold for reacting to the cross-reactive self-antigens is effectively lower. This allows the T cells activated by a pathogenic peptide that are cross-reactive to a self-antigen to be added to the pool of T cells that are usually robustly activated by this self-antigen. This can result in exceeding the quorum threshold, thus resulting in a functional autoimmune response. Pathogen-derived peptides that satisfy these conditions are rare. Persistent or severe infection leads to increasing numbers of pathogen-derived peptides being targeted, which increases the probability of sampling such peptides and overcoming the quorum threshold for responses to certain self-peptides. Local inflammatory conditions and tissue damage during persistent viral infections can increase the presentation of different self-antigens. This increases the probability of finding pairs of pathogen-derived and self-antigens with cross-reactive T cells that can exceed the quorum threshold for the self-antigens, resulting in autoimmunity. Our model provides an explanation for why individuals who are persistently infected with viruses like EBV and EV develop the corresponding autoimmune disease only with some probability, as well as the latency of pathogenesis.

We first develop and employ a computational model of T cell selection in the thymus to show that appropriately set values of positive and negative selection thresholds result in mature repertoires that contain autoreactive T cells while maintaining tolerance to self-antigens with high probability because the quorum threshold is not exceeded by T cells in response to self-antigens alone. We then use our computational model to study the response of such T cell repertoires to infection. Consistent with data [63, 58], we assume that a persistent or severe infection results in the presentation of multiple pathogen-derived peptides either sequentially or at the same time. Our computational results show that the probability of triggering autoimmunity increases monotonically with increasing numbers of pathogen-derived pMHC molecules that are targeted by T cells. Further analyses of our computational results show that rare pathogen-derived peptides that could trigger activation of many T cells may play a significant role in mediating autoimmunity upon persistent viral infection. Importantly, we propose specific experiments in mouse models that could directly test the importance of how collective effects and molecular mimicry are intertwined to break tolerance upon persistent viral infections.

### Development of a simple computational model of thymic selection and immune responses of the mature T cell repertoire

In this section, we first describe our computational model for development of the mature T cell repertoire in the thymus. We then utilize our model to characterize the response of the thus generated mature repertoire to infection (i.e., pathogen-derived pMHC molecules), and the impact of infection on the potential emergence of autoimmunity.

#### Model for development of the T cell repertoire

During development, a thymocyte encounters a number of self-pMHC complexes displayed on cells in the thymus and undergoes positive and negative selection. These processes occur in stages in different compartments of the thymus; for example, negative selection occurs in two different stages. These details of stages of selection are not incorporated in our simple model. Each interaction of a thymocyte’s TCR with a self-pMHC corresponds to a different binding free energy. We use a convention in which lower binding free energy corresponds to higher affinity. To successfully mature and exit the thymus, the binding free energy of a thymocyte’s TCR to at least one of these pMHCs must be lower than E_p_ (positive selection). The binding free energy must also not be lower than a more negative value, E_n_, for interactions with any of the encountered self-pMHC as otherwise the thymocyte is negatively selected. This implies that the minimum value of the binding free energy of a thymocyte with the encountered self-pMHC molecules must lie between E_p_ and E_n_. Therefore, our model for the development of a mature T cell repertoire involves generating a panel of self-pMHC molecules and thymocytes (each expressing a TCR), calculating the binding free energies of every TCR with each encountered self-pMHC molecule, and checking whether the minimum value lies between E_p_ and E_n_. Thymocytes with TCRs that satisfy this condition are added to the mature T cell repertoire [Fig. 3A]. To carry out this procedure, we need to define a model for TCRs expressed on thymocytes and self-pMHC molecules, and their binding free energies. This model will also be used to study the response of the mature repertoire to pathogen-derived pMHC molecules upon infection.

**Figure 3:**
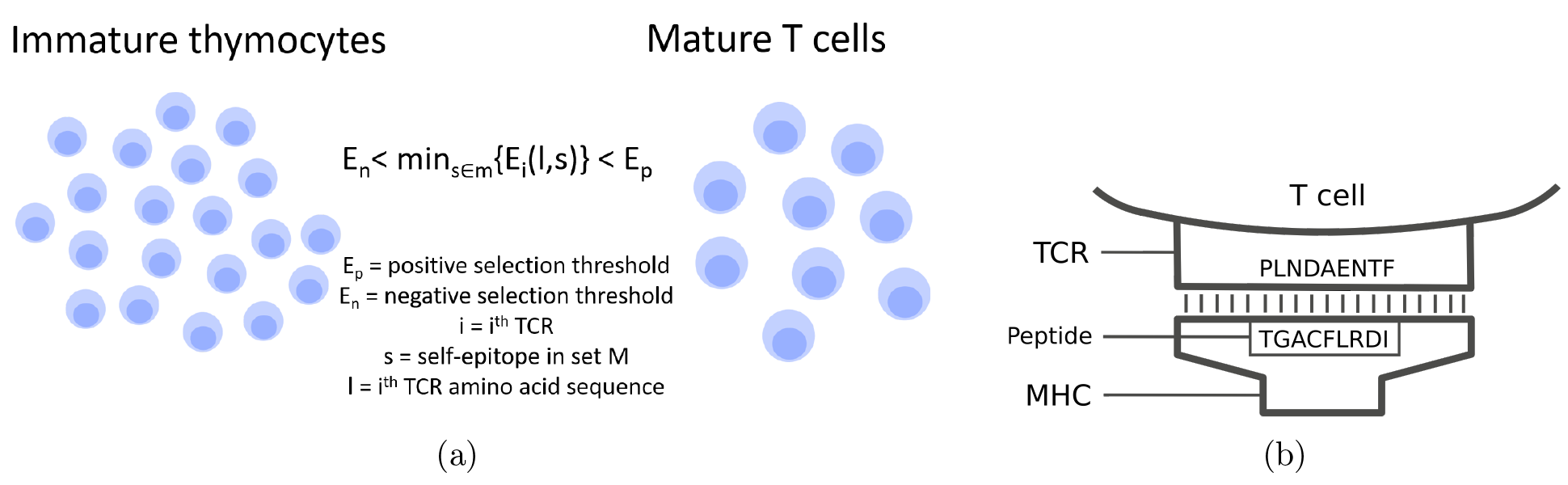
Depiction of thymic selection processes and model for TCR-pMHC binding free energy: (a) Modeling thymic selection: only thymocytes with a minimum binding free energy between E_p_ and E_n_ go on to mature. (b) Model for TCR-pMHC binding free energy: The variable residues of the TCR that interact with the peptide in the pMHC complex are represented by strings of amino acids. Addition of pairwise interactions of amino acid along the two strings provides the total binding free energy E_i_ between a TCR-pMHC pair.

Our model for TCRs and pMHC molecules and their binding free energies follows the framework of Ref. [19], in which TCR and pMHC molecules are represented as strings of amino acids. TCRs interact with the peptide and the MHC. Past work using a similar model has argued that interaction free energies with the MHC are drawn from a relatively narrow distribution, and including these interactions do not change the qualitative results pertinent to the characteristics of the mature repertoire. Therefore, we do not explicitly include MHC, and represent only the peptides and the peptide contact residues of TCRs as strings of amino acids [Fig. 3B]. We assume that each peptide contact residue on a TCR interacts only with a corresponding site on a peptide. Previous work with a similar model that included more complex interaction patterns between these sites showed that most qualitative results are not altered by considering this complexity [75]. For a given TCR-pMHC pair, the interaction free energy is calculated as:

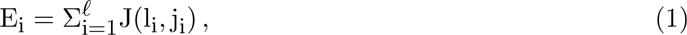

where *ℓ* is the length of the variable regions of the TCRs and pMHCs as seen in Figure 3B, and J(l_i_, j_i_) is the interaction free energy between the amino acid at the i-th peptide contact residue of the TCR (l_i_) and the peptide (j_i_). For instance, in Figure 3B, the interaction free energy between the first peptide contact residue of the TCR and peptide is that of Proline (P) and Threonine (T), and the total interaction free energy is the summation of the interaction free energies for each pair of amino acids between the TCR (PLNDAENTF) and the peptide (TGACFLRDI). Here, we took *ℓ* to be 9, the approximate length of MHC class I binding peptides. However, the model is formulated to be MHC class agnostic and could apply to either class. Moreover, reducing the number of contact residues, as they are typically fewer [Section Experimentally testable predictions of the model][22] should not change our qualitative results. The interaction free energy between a pair of amino acids in the TCR and peptide is calculated using the Miyazawa-Jernigan (MJ) matrix in units of k_B_T [76].

This simple model of TCR-pMHC interactions has been used previously to study thymic development. One prediction that emerged is that positive and negative selection tune the statistical distribution of the biochemical properties of the amino acids in the peptide contact residues of the TCRs on mature T cells. The peptide contact residues of TCRs in the mature repertoire were predicted to be statistically enriched in amino acids that interact moderately with other amino acids [19, 20, 21]. Attesting to the utility of the model, this prediction was tested positively in mouse models [22]. Moderately hydrophobic amino acids were found to be statistically enriched in the key peptide contact residues of TCRs in mature T cells, and this property of the mature T cell repertoire was dependent on normal negative selection. Hydrophobicity is a natural measure of the strength of interactions at an interface, such as for TCR-pMHC binding.

Although statistical models for VDJ recombination are available [3, 77, 78], for simplicity, the sequences of peptide contact residues of the TCRs of thymocytes were generated using just the amino acid frequencies that characterize the human proteome [Supp. 1]; 10^6^ unique immature thymocytes were generated. As we will note later, because T cell clonality is not considered in our analyses, our results represent a conservative estimate of the probability of triggering autoimmunity upon persistent viral infection. To generate sequences of self-peptides, we note that not all peptides can be properly presented by all MHCs. To construct a library of self-peptides to present to thymocytes, we screened self-peptides for presentation by human MHC alleles (Human leukocyte Antigen or HLA) using NetMHCpan [79]. This bioinformatics tool is trained on measured peptideHLA-I interactions and predicts binding with 95% accuracy for each HLA allele. We focused on HLA-A*01:01, HLA-A*02:01, HLA-B*08:01 and HLA-B*07:02, the most commonly found HLAs in the human population. NetMHCpan frameshifts the human proteome from UniProt [80, 81] and determines whether each peptide of length 9 thus generated binds to a specified HLA. We then took the sequences of peptides predicted to bind to a particular HLA and compared their binding affinities to the distribution of affinities calculated for a set of randomly chosen peptides that have been found to be presented on HLA in human samples; only those predicted binders with HLA binding affinities in the top 2% of the latter affinity distribution were chosen to be among our set of self-pMHC molecules. Thus, we generated 10,000 self-peptides (N) with equal numbers of peptides that bind to HLA-A*01:01, HLA-A*02:01, HLA-B*08:01 and HLA-B*07:02.

To generate the mature T cell repertoire as described above, we expose each TCR generated as described above to 8,000 self-peptides chosen randomly from our set of 10,000 self-peptides. We generate several T cell repertoires to obtain our results (see later), and each time a different random set of 8,000 self-peptides is chosen. We need to pick appropriate values for E_p_ and E_n_ to obtain the results of selection. To do so, we simulated thymic selection with different choices of E_n_ and the gap Δ = E_p_ – E_n_. For each value of E_n_, the value for the gap Δ was chosen such that 4% of thymocytes survived selection, as observed in experiments in mouse models [82]. As there are typically two values of E_n_ that satisfy this requirement [Supp. 2], the value for which negative selection is limiting was chosen [20]. This is consistent with experiments in mouse models [22] that show that defects in negative selection change important statistical characteristics of the mature T cell repertoire, such as the frequency of hydrophobic amino acids that make up TCR peptide contact residues. Our results are not sensitive to the exact value of E_n_ (as long as negative selection is limiting) [Supp. 2]; we used E_n_ = –38k_B_T for the results described in the main text.

### Modeling infection and its impact on autoimmunity

During an acute infection, only a few pathogen-derived peptides are immunodominantly targeted by T cells [83]. During persistent or severe infection, either as time ensues or at the same time, many more peptides can be targeted by T cells. We thus consider infection with different numbers of pathogen-derived peptides (N_f_), and ask whether the T cell response to these peptides can also trigger a functional autoimmune response. The chosen values of N_f_ range from 0 to 10, with higher values of N_f_ representing a persistent or severe infection. Taking *Listeria monocytogenes* as a model pathogen and its proteome from UniProt [84], we used NetMHCpan to generate 40,000 peptides that can be presented by the same human HLA alleles used in thymic selection of the mature T cell repertoire.

Figure 4 outlines the steps in our method for modeling infection. We first generate a particular realization of a mature naive T cell repertoire using the method in the preceding section. Then, we randomly pick one of the 40,000 Listeria peptides, as well as N_T_ self-peptides from our panel of 10,000, assuming that both sets can be presented by HLAs in the infected tissue. Next, we calculate the binding free energies of the mature T cells in the repertoire with the pathogen-derived peptide. If this free energy is lower (higher affinity) than the activation threshold, taken to be E_n_, we count the T cell as being activated. Thus, we determine how many T cells are activated by the chosen pathogen-derived peptide. We chose a sharp threshold of activation because in individual T cells the membrane-proximal signaling network exhibits a digital, all-or-nothing response [85, 86, 87]. Setting the activation threshold to equal E_n_ ensures that mature T cells that survive selection are not activated by the self-peptides they encountered during thymic selection [23]. If the number of T cells activated by the pathogen-derived peptide is below the quorum number Q, then this event is counted as non-infectious, and also does not trigger autoimmunity. If the number of T cells activated by the pathogen-derived peptide exceeds Q, we study the impact of this functional immune response on triggering autoimmunity. Following indications from experiment [45, 51], we choose the quorum number to be Q = 10. Note that this value is likely the average value of Q, a point that we will elaborate upon later.

**Figure 4:**
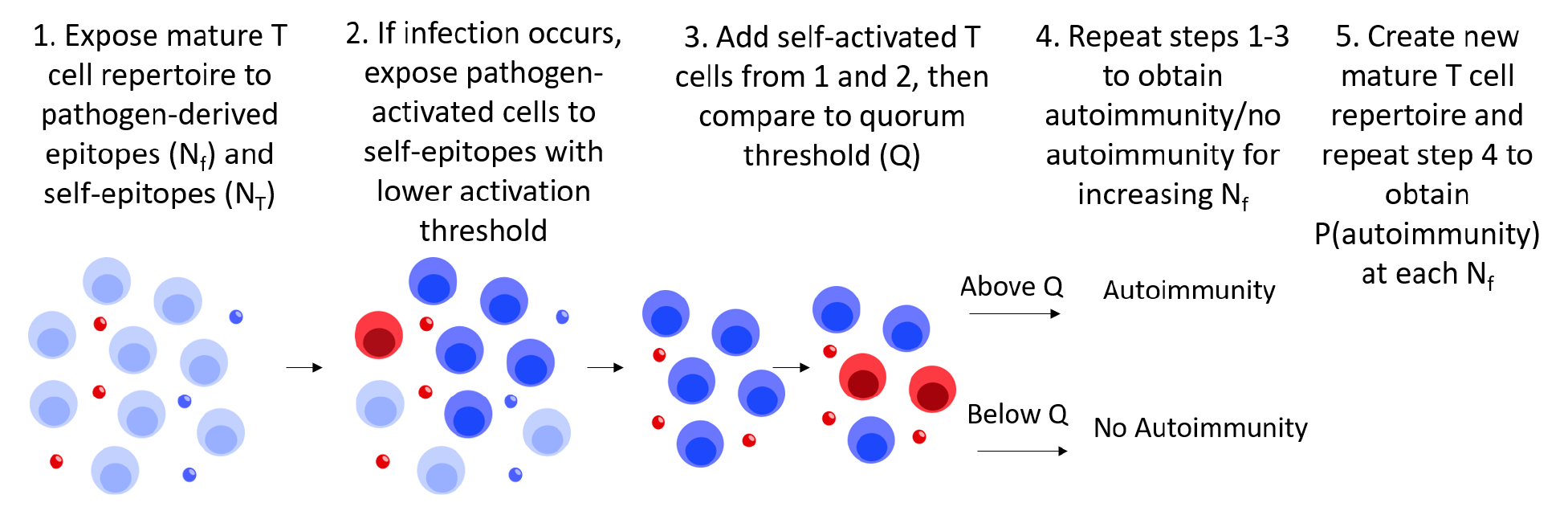
Steps in modeling infection: We follow the steps shown and described in the text for increasing numbers of pathogen-derived peptides (or epitopes), N_f_, and the same number, N_T_, of self-peptides to calculate the probability of a functional autoimmune response.

**Figure 5:**
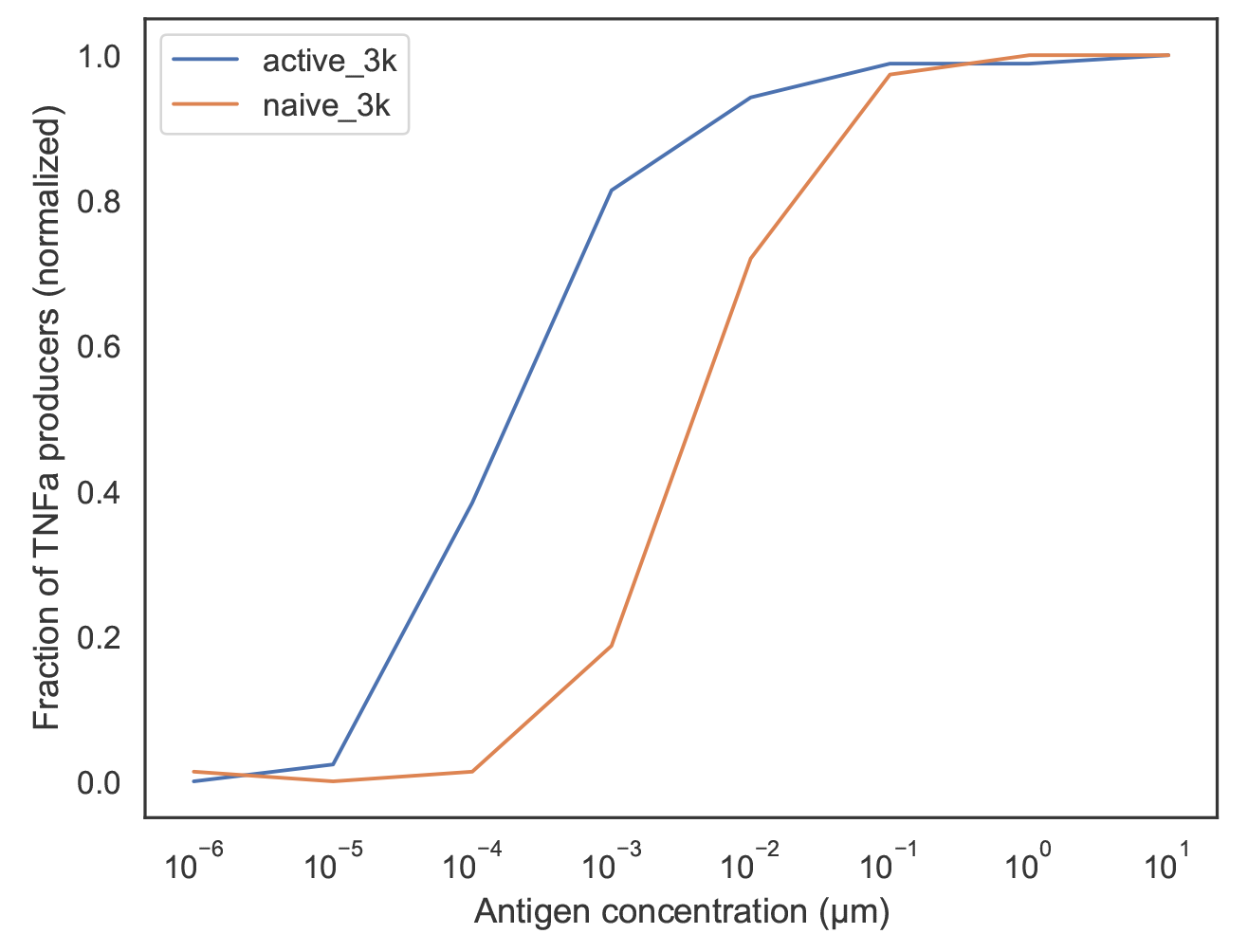
Graph of antigen concentration and % of T cell population that are TNFα producers when exposed to the 3k antigen. The red and blue curves correspond to the responses of naive and previously activated T cells, respectively.

When there is a functional response to the infection, we take each T cell activated by the pathogen-derived peptide and calculate its binding free energy with the N_T_ self-peptides presented in the tissue. The goal is to determine if these activated T cells are cross-reactive to the self-peptides in the tissue. Checking for cross-reactivity is consistent with the concept of molecular mimicry. To determine whether or not an activated T cell is crossreactive to each self-peptide in the tissue, we account for the fact that activated T cells have a lower threshold for re-activation. Therefore, for T cells previously activated by a pathogen-derived peptide, the binding free energy threshold for reactivation upon interactions of its TCR with the self-pMHC molecules is taken to be higher than E_n_. The value of this lower binding free energy, E_weak_, is chosen based on our experimental data [Fig. 5]. We measured the response of a population of B3K506 TCR Tg T cells with receptors specific for an antigen labelled 3K. The response of both naive and activated T cells were measured using titrating concentrations of peptide presented by dendritic cells. The fraction of responding T cells was determined by measuring whether they produce the cytokine TNFα [Fig. 5]. Another example of this type of data for a different peptide (P5R) is shown in Supp. 3.

We used the response curves in Figure 5 to estimate E_n_–E_weak_. While individual T cell activation is digital, activation at the population level has a sigmoidal shape [85]. However, a typical T cell activation threshold can still be inferred from the population response curve [23] by noting that the experimental curves resemble standard binding fractions as a function of chemical potential μ and binding free energy E_B_, given by

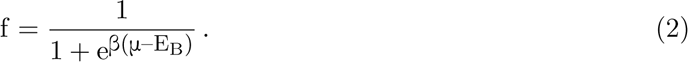

The chemical potential at a response fraction of 1/2 corresponds to the binding free energy. Since in a dilute solution concentration (c) is proportional to e^βμ^, the difference in concentrations at a response fraction of 1/2 for naive and activated T cells in Figure 5 can be related to differences in binding free energies by

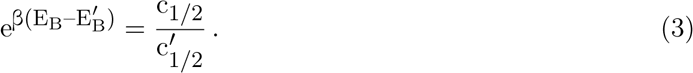

By comparing the concentrations at a response fraction of 1/2 in Figure 5, we estimate E_weak_ – E_n_ *≈* 3k_B_T, and using E_n_ = –38k_B_T we obtain E_weak_ = –35k_B_T.

Using this lower activation threshold, we tested whether any of the pathogen-activated T cells are also activated by the panel of N_T_ self-peptides presented in the tissue. Let us denote the total number of crossreactive T cells activated by all N_T_ self-peptides by N*^′^*. If the sum of T cells activated by these N_T_ self-peptides before infection and N*^′^*_self_ exceeds the quorum threshold, this event is counted as one that triggers autoimmunity upon infection. This procedure is repeated if additional pathogenic peptides are targeted by T cells during infection. If we used peptide A for the one pathogen-derived peptide (N_f_ = 1), for N_f_ = 2 we add a second peptide, B. If B also activates a larger number of T cells than Q, these T cells are also tested for crossreactivity with the N_T_ self-peptides in the tissue. The crossreactive ones are added to the pool of T cells that could exceed the quorum threshold against the self-antigens, and if the quorum threshold is exceeded this counts as an autoimmune event if the two pathogen-derived peptides, A and B, are both targeted. This procedure is continued sequentially until a desired value of N_f_ is reached. To obtain statistically meaningful results for the probability of triggering autoimmunity for each value of N_f_, we repeat the above calculations 10,000 times, with each trial carried out with new choices of the N_f_ pathogen-derived peptides and the N_T_ self-peptides displayed in the tissue, but with the same mature T cell repertoire. The probabilities thus obtained represent the average chance that autoimmunity will be triggered in a particular individual with a specific T cell repertoire for each value of N_f_ . Finally, all the calculations described above are carried out for 30 different realizations of the mature T cell repertoire (or individuals), and the average value for triggering autoimmunity is reported for each value of N_f_ . Table 1 summarizes the parameter values used in our simulations.

**Table 1:** Parameters used in our simulations.

| Parameter | Value |
| --- | --- |
| Number of immature thymocyte | 1,000,000 |
| Fraction of surviving thymocytes | 0.04 |
| Number of surviving thymocytes | 40,000 |
| $E_n$ | -38k<sub>B</sub>T |
| Number of self-peptides presented during selection (M) | 8,000 |
| Number of self-peptides in total (N) | 10,000 |
| Number of pathogenic peptides in total | 40,000 |
| Number of self-peptides presented in tissue ($N_T$) | 10 |
| $E_{\text{weak}}$ | -35 k<sub>B</sub>T |
| Quorum threshold | 10 |
| Number of trials | 10,000 |

## Results

As noted earlier, consistent with data [63, 58], we assume that a persistent or severe infection results in the presentation of multiple pathogen-derived peptides either sequentially or at the same time. Thus, the variation of the probability of triggering autoimmunity as a function of the number of pathogen-derived peptides targeted in a tissue (N_f_) should shed light on differences in the chance of triggering autoimmunity upon persistent infection or severe infections and more usual infections, with severe or persistent infections corresponding to larger values of N_f_ . In the subsections to follow, we will first describe the results of our simulations and then elaborate the mechanistic reasons that underlie our results.

### The probability of triggering autoimmunity grows with the number of pathogen-derived peptides targeted by T cells during infection

As depicted in Figure 6A, simulations show that increasing the number of pathogen-derived peptides targeted by T cells also increases the probability of triggering autoimmunity. The increased chance of autoimmunity with N_f_ provides an explanation for why persistent or severe infections can increase the chance of developing autoimmune conditions. As the probability of developing autoimmunity is not large, not everyone with a persistent infection develops autoimmunity. Our results also explain why there is a lag time between the establishment of persistent infection and the onset of autoimmunity. As time ensues, increasing numbers of pathogen-derived peptides are presented and targeted by T cells, and the chance of triggering autoimmunity grows as per the results reported in Figure 6.

**Figure 6:**
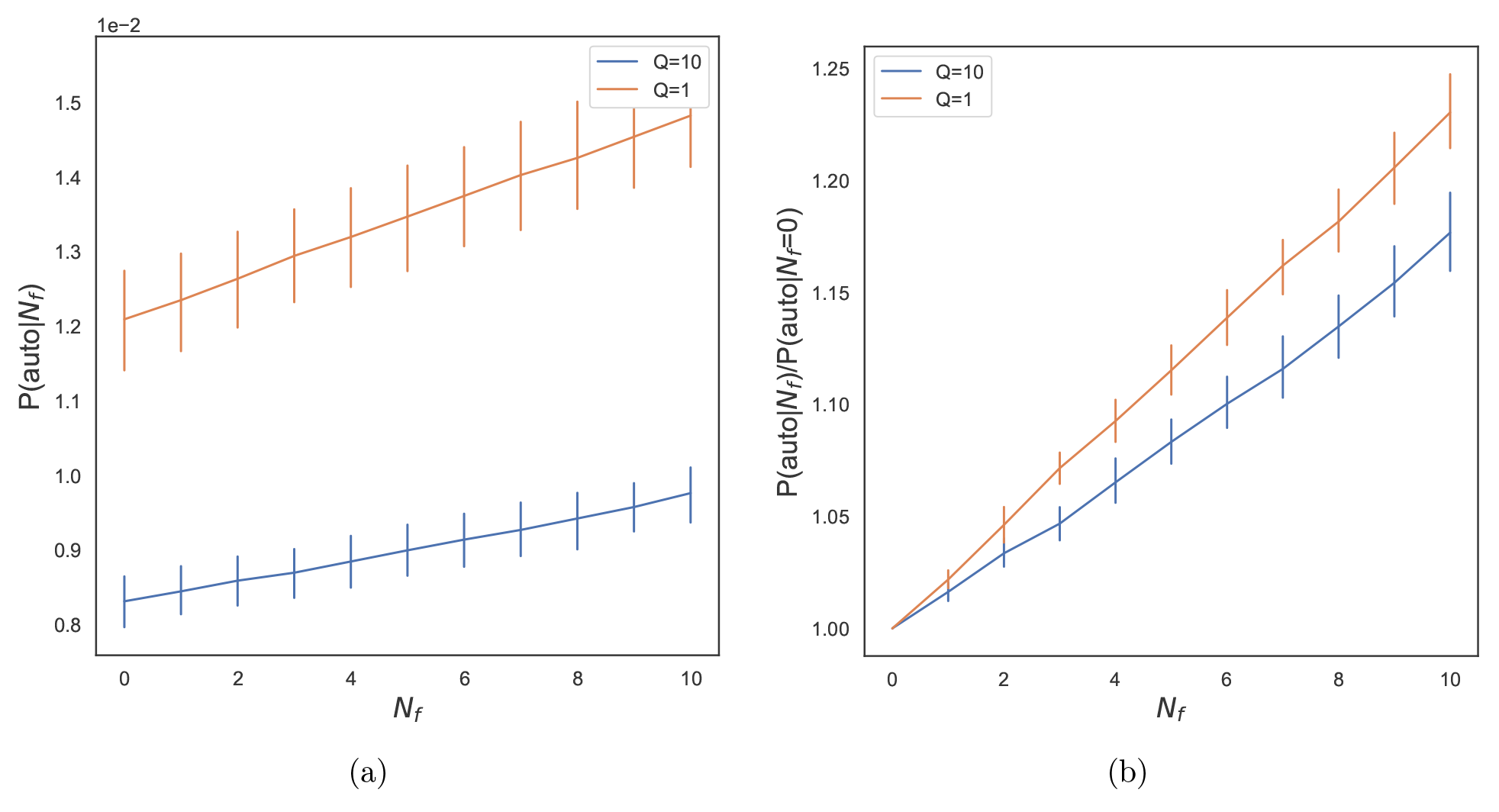
(a) Probability of triggering autoimmunity, P(auto|N_f_), as a function of the number of pathogen-derived peptides targeted for two different values of Q, Q = 1 (no collective effects) and Q = 10 (collective effects as per quorum model). (b) Probability of triggering autoimmunity, P(auto|N_f_) normalized by the probability of triggering autoimmunity in the case of no infection, P(auto|N_f_ =0). Parameters used are as shown in Table 1. The error bars were calculated based on the number of simulations noted in Figure 4.

Our results also explain why only certain viral infections trigger particular autoimmune conditions. A key component of our model is that T cells activated by pathogen-derived peptides are weakly cross-reactive to self-peptides presented in the same tissue. Such crossreactivity or molecular mimicry is only possible for certain self-antigens and pathogen-derived peptides, such as the observed crossreactivity between EBV and myelin-derived peptides in the case of MS [66, 88, 67] and microbial and self-peptides in the case of AS [67]. However, such crossreactivity exists regardless of the collective effects inherent in the quorum model. Figures 6A and 6B show the influence of collective effects as we compare what happens when Q = 1 (no collective effects) and Q = 10. Increasing the quorum number, Q, not only has the desired effect of reducing the intrinsic probability of autoimmunity due to circulating autoreactive T cells in the absence of infection (N_f_ = 0), but also decreases the relative chance of triggering autoimmunity upon viral infections [Fig. 6A]. This is because two conditions that depend on collective effects must be satisfied to trigger autoimmunity. First, the number of T cells activated by a pathogen-derived peptide must exceed the quorum threshold and then the ones among these T cells that are crossreactive to self-antigens in the same tissue must contribute to exceeding the quorum threshold with the self-antigens. In some instances, it is possible that a pathogen-derived peptide can trigger a sufficiently large number of T cells that the ones among these that are proliferating and cross-reactive to a self-antigen are the major contributors to exceeding the quorum threshold with the self-antigen and triggering autoimmunity. We will discuss this point more fully in the next section. As noted earlier, we do not consider multiple clones of the same T cell. Including clonal T cell populations while still using exactly our model would make the probability of triggering autoimmunity higher compared to that reported in Figure 6.

The number of self-peptides that T cells encounter in the thymus can vary from person to person and is different for different T cells. Also, experimental results indicate that a HLA allele associated with diabetes binds self-peptides in a less stable way [89], suggesting that T cells restricted by this HLA allele are likely to encounter fewer self-peptides during development. We investigated whether the collective effects embodied in the quorum model can make the immune system more robust by inhibiting autoimmune responses upon persistent or severe infections even in the face of such variations. To address this question, we carried out simulations with different values for the average number of self-peptides T cells encounter in the thymus (M) and calculated the change in probability of triggering autoimmunity upon persistent viral infection (N_f_ = 10) for different values of Q. Figure 7A shows the results of simulations for M = 8, 000 (as in Fig. 6) and a lower value, M = 6, 500. The increase in the probability of autoimmunity being triggered if M = 6, 500 is shown for various values of the quorum threshold, Q. As the value of the quorum threshold increases, the increase in the probability of autoimmunity decreases. Thus, the collective effects embodied in the quorum model makes the immune system more robust to inter-person variations in thymic development by lowering the change in the probability of triggering autoimmunity upon persistent infections due to such variations. We also find similar results if N_T_, the number of self-peptides presented in a tissue, increases from 10 to 20 [Fig. 7B]. Our results in Figure 7 indicate that the majority of benefits of the robustness due to collective effects is realized at Q *∼* 10. While our calculations are not meant to be quantitatively accurate, it is interesting to note that this value is consistent with experimental observations that the number of proximal activated T cells required for proliferation and differentiation is of the same order. [45][51].

**Figure 7:**
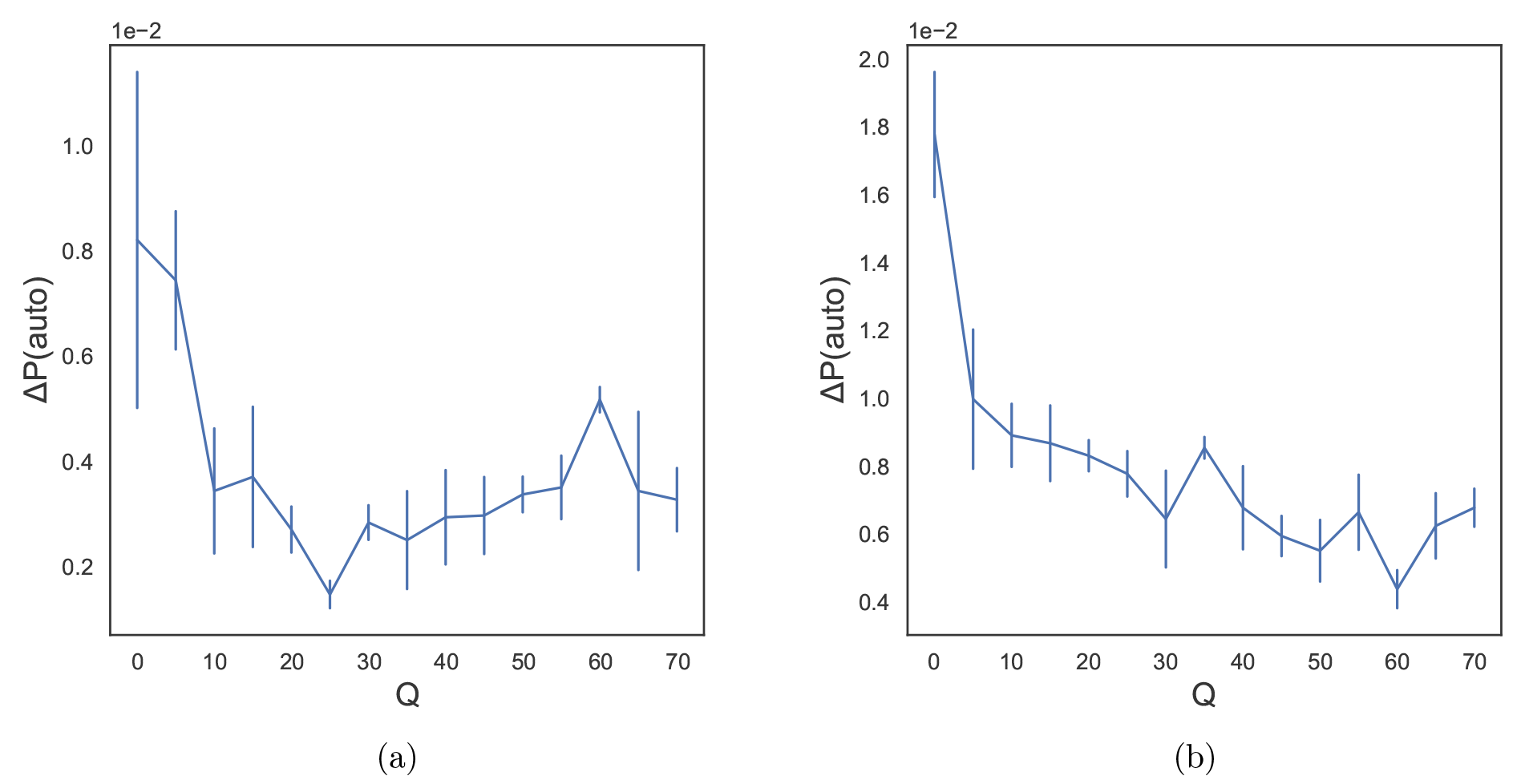
Collective effects in the quorum model confer robustness against autoimmunity. All simulations unless otherwise denoted were performed with the same parameters as in Table 1. (a) Simulations were performed with M=8,000 and M=6,500 at increasing values of Q at N_f_ = 10. The calculated change in the probability of triggering autoimmunity, P(auto), is graphed as a function of Q. (b) Simulations were performed with N_T_ = 10 and N_T_=20 for increasing values of Q at N_f_ = 10. The calculated change in P(auto) is graphed as a function of Q.

The effector functions of activated T cells that target infected cells can lead to damage to healthy tissue and the release of otherwise sequestered self-antigens into the local tissue, thus increasing the presentation of self-antigens [90, 91]. We wondered whether the collective effects inherent to the quorum model might also inhibit the increased chance of autoimmunity being triggered due to enhanced presentation of self-antigens. Figure 7 shows results of our simulations for the increase in probability of triggering autoimmunity as a function of Q when the number of self-antigens presented in the tissue (N_T_) increases from N_T_ = 10 to N_T_ = 20. We find that increasing values of Q leads to a smaller increase in the risk of autoimmunity given an increase in self-antigen presentation. These results suggest that autoimmunity being triggered by epitope spreading and bystander activation [56] is reduced due to collective effects [Fig. 7]. Again, the influence of collective effects in suppressing autoimmunity upon strong perturbations diminishes once the quorum threshold is sufficiently high [Fig. 7]. It is important to note, however, that increasing N_T_ leads to a monotonically increasing risk of autoimmunity in spite of the collective effects embodied in the quorum model [Supp. 4]. This result shows that, in spite of the robustness conferred by the quorum model, substantial changes in the number of self-peptides presented in a tissue enhances the probability of triggering autoimmunity. So, if increased inflammation and tissue damage result in presentation of new self-antigens, autoimmunity would be more likely to be triggered upon infection.

The results described in this section suggest that collective effects embodied in the quorum model not only confer robustness to the immune system from the standpoint of discriminating between self and pathogen-derived antigens, but also serve to inhibit autoimmunity in spite of inter-person variations in thymic development and when strongly challenged such as upon severe or persistent infection.

### The importance of rare, highly reactive pathogen-derived peptides for triggering autoimmunity

In order to obtain additional mechanistic insights, we first attempted to analyze the simulation results reported above in terms of a simple probabilistic model. The model is framed in terms of the following probabilities:

- q, the probability that a pathogen-derived peptide activates a T cell. Thymic selection against M peptides leads to q *∼* 1/M [37].
- q_s_, the probability that a self-peptide activates a T cell. If M out of N possible self-peptides are seen in the thymus, q_s_ = q(1 – M/N).
- q_w_ = αq_s_, the probability that a pathogen-activated T cell can be reactivated by a weakly crossreactive self-peptide. Cross-reactivity is thus encoded in the parameter α *>* 1 that can be deduced from Figure 5.

An important consideration is how many of the T cells patrolling a tissue are on average activated when a self or pathogen-derived peptide is presented. These numbers are estimated using the parameters of Table 1, as n = Tq *≈* 10^6^ *×* 0.04 *×* (8, 000)^−1^ *≈* 5 for pathogen-derived peptides, and n_s_ *≈* n(1 – 8, 000/10, 000) *≈* 1 for self-peptides. *Assuming that T cell activation events are independent* leads to a Poisson distribution for the probability of activating m T cells, and a probability of exceeding the quorum number Q of

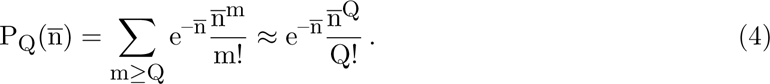

The last approximation is correct as long as quorum number Q *>* n which holds for the choice of Q = 10. This leads to an estimate of a roughly 2% probability for the quorum threshold being exceeded by a pathogen-derived peptide (on average), and a probability of 10^−7^ for a self-peptide (obtained as P_Q_(n_s_)).

However, the base value of autoimmunity in the absence of infection (N_f_ = 0) in Figure 6A is orders of magnitude higher than the 10^−7^ predicted by Eq. 4 for P_Q_(n_s_). This discrepancy can be traced back to the Poisson distribution not capturing the characteristics of the T cell repertoire. Figure 8A shows results of our simulations for the distribution of the number of T cells in the mature repertoire that are activated upon being challenged by our set of N_f_ = 40, 000 pathogen-derived peptides and N_s_ = 10, 000 self-peptides. These simulations were carried out with many mature T cell repertoires and choices of the self and pathogen-derived peptides. For both self and pathogen-derived antigens, the distribution of the number of activated T cells is very different from a Poisson distribution and is characterized by a long tail. That is, rare peptides can activate a large number of T cells. For example, a peptide composed of highly hydrophobic amino acids would be able to activate many T cells with TCRs containing similar moderately or poorly hydrophobic peptide contact residues. Thus, the activation of these T cells would be correlated and not independent, and the assumption of a Poisson distribution in Eq. 4 would be incorrect. This characteristic of rare peptides being able to activate many T cells can be illustrated with a much simpler model of TCR-peptide interactions than the string model.

**Figure 8:**
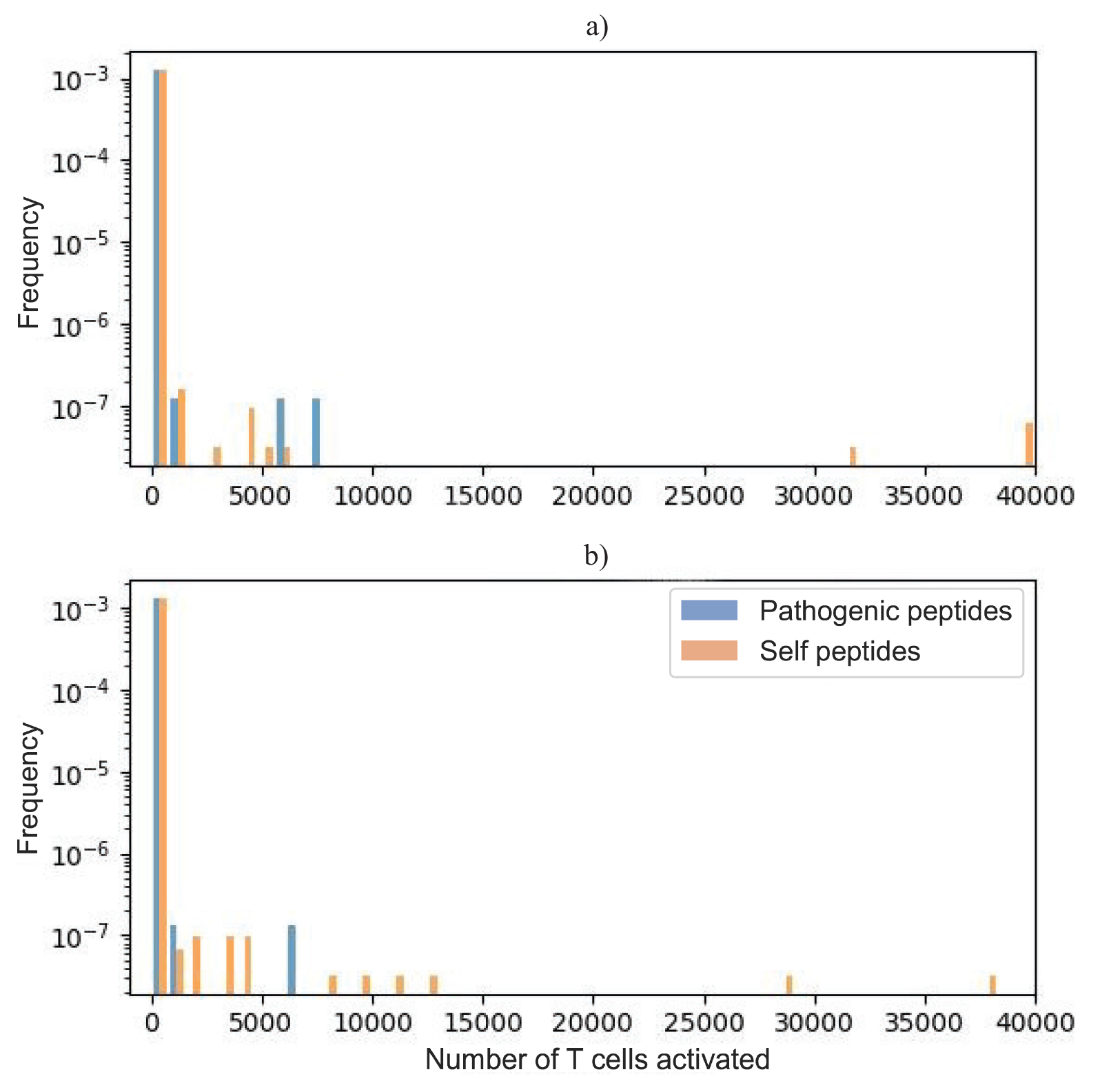
(a) Distribution of number of T cells activated by pathogen-derived and self-peptides as obtained from our simulations. Each simulation was conducted with M = 8,000 and 40,000 pathogen-derived peptides, and 10,000 self-peptides. 9,987 self-peptides activated 0 T cells, 5 self-peptides activated between 1 and 10, and 8 self-peptides activated 10 or more. 39,901 pathogen-derived peptides activated 0 T cells, 43 activated between 1 and 10, and 56 activated 10 or more. (b) Distribution of number of T cells activated by pathogen-derived and self-peptides as obtained from the simpler coarse-grained model described in the text. 9,995 self-peptides activated 0 T cells, 1 self-peptide activated between 1 and 10, and 4 self-peptides activated 10 or more. 39,983 pathogen-derived peptides activated 0 T cells, 4 activated between 1 and 10, and 13 activated 10 or more.

For a given peptide sequence j, in the string model we can construct a reactivity parameter β_j_ as the mean of the strength of its interactions with all possible TCR sequences. The ensemble of possible peptides can be characterized by the probability distribution p(β) of reactivity. We can construct a similar distribution p(α) describing the mean strength of interactions of pre-selection thymocyte sequences (each sequence, i corresponds to α_i_). To illustrate the role of reactivity, we consider a simplified model in which thymocyte sequences α_i_ and β_j_ are taken from the same Gaussian distribution, and their binding free energy is computed simply as

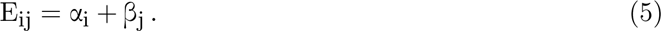

Positive and negative selection prunes the initial TCR ensemble to create a post-selection set of TCRs characterized by a probability distribution, p(α_i_), which is concentrated in the range α_min_ *<* α_i_ *<* α_max_ *<* 0 [Supp. 5]. Possible values of β_j_, however, are not similarly restricted, and it is possible (within the model) to encounter rare self-peptides β_j_ such that β_j_ + α_min_ *<* E_n_; such peptides would then activate many T cells. We carried out calculations with this “coarse-grained” model with parameters close to that used in simulations with the “string” model [Supp. 5]. The resulting distributions for number of activated T cells are presented in Figure 8B. The distribution of T cell reactivity to peptides is similar to that obtained using the string model, including a tail of rare highly reactive peptides that activate many T cells that can overcome the quorum threshold. The chance of sampling such rare peptides from this tail of the distribution grows with the number of pathogen-derived peptides that are targeted as is the case during persistent or severe infections, but not usual acute infections. Such a tail of highly immunogenic antigens has also been observed in the context of SARS-CoV-2 infection. Out of a panel of HLA-I SARS-CoV-2 epitopes, 122 were considered immunogenic [92]. This is approximately 0.4% of total possible HLA-I restricted epitopes, which is comparable to the percentage of pathogen-derived peptides above the quorum number in our simulations (roughly 0.14%).

We emphasize that our goal is not quantitative recapitulation of known experimental facts. Our results are not quantitatively accurate as they are derived from simple models that aim to uncover new mechanistic insights that lead to experimentally testable predictions (see next section). For example, Figure 8 suggests that rare self-peptides would trigger autoimmunity by overcoming the quorum threshold, and so over long times, they should cause autoimmunity in everyone. But, this result is due to a limitation of our simple model. In our model the quorum threshold, Q, is fixed. In reality, T cells that bind to self-peptides in the thymus with long half-lives are likely to differentiate into Tregs [93]. Therefore, many more conventional effector T cells would have to be activated to beat the suppressive effects of more numerous Tregs that are activated by the same rare self-peptides; i.e., the quorum threshold is not a fixed number, but depends upon the T cell reactivity of the self-peptides. We also note that the Mirazawa-Jerningan model we have used for interactions between amino acids [76] is known to overemphasize the strength of hydrophobic interactions, which would result in an overestimate of the strength of TCR-pMHC interactions.

The main insight from the results presented in this section is that rare crossreactive self and pathogen-derived peptides may work together to trigger autoimmunity by exceeding the pertinent quorum thresholds. Persistent or severe infection makes the chance of sampling these peptides more likely.

## Experimentally testable predictions of the model

It has been proposed that pathogen-derived peptides with homologs in the host proteome should be associated with higher likelihoods of triggering autoimmunity [94]. But, this notion does not account for the importance of collective effects embodied in the quorum model and the results we have described above. We propose immunizing mice with increasing numbers of pathogen-derived peptides that are homologous to peptides derived from the mouse proteome. The prediction of our model is that, as the number of such pathogen-derived peptide immunogens introduced into the animals increases, the chance of triggering autoimmunity should also increase.

To experimentally test this prediction, we have designed immunogens using the following approach. We propose immunizing C57BL/6 (B6) mice that express the I-Ab MHC molecule on their APCs. We used NetMHCIIPan to find 15-mer I-Ab binding *mycobacterium tuberculosis* peptides and peptides from the mouse proteome, respectively [79, 95, 96]. Then, following past work [94], we identified mouse homologs for each of the tuberculosis peptides by identifying the mouse peptides with 9-mer cores that shared the same amino acids in the 2,3,5,7, and 8 positions with the tuberculosis peptide’s 9-mer core. The number of mouse homologs per tuberculosis peptide ranged from 0 to 3, 000 [Supp. 6]. These peptides that have similarity to the I-Ab binding mouse peptides are promising candidates for our proposed experiment, as we expect them to be more likely to activate cross-reactive T cells than peptides with low similarity to self-peptides. However, the pathogen-derived peptides should also be chosen to be different enough from self-peptides such that thymic selection would not have negatively selected against T cells that could recognize them with high probability. Based on these considerations, we chose tuberculosis peptides that are in neither of the tails of the similarity distribution (i.e., not too similar and not too distinct from self-peptides). We thus selected tuberculosis peptides that have 17 mouse homologs, which lies approximately at the mean of the similarity distribution shown in Supp. 6. These 122 peptides are listed in the Table provided in Supp. 6. Because we expect the rare peptides that activate many T cells to have more hydrophobic amino acids [22], we have also indicated the peptides in our list of homologs that have the most hydrophobic amino acids. Hydrophobicity was determined using multiple methods (KyteDoolittle [97], Hopp-Woods [98], Eisenburg [99], Rose [100], Engelman [101], and Wimley-White [102]) and essentially the same peptides were identified as being comprised of the most hydrophobic amino acids.

We suggest a protocol wherein mice are immunized with randomly picked tuberculosis peptides from our list. We predict that immunization with an increasing number of these peptides is more likely to trigger autoimmunity. We also anticipate that autoimmunity is more likely to be triggered upon immunizing with peptides from our list that we have identified as having more hydrophobic amino acids.

## Discussion

In recent years, based on theoretical modeling, experiments in mice and in vitro, and analyses of sequences of T cell repertoires, evidence for the importance of collective effects in mediating T cell proliferation, differentiation and a functional response has accumulated [37, 44, 45, 10, 52]. These data suggest that a threshold number (or density) of T cells must be activated in a tissue in order for activated T cells to proliferate and mount an immune response. This threshold number, or quorum, is necessary to beat out the suppressive effects of peripheral tolerance mechanisms mediated by Tregs and perhaps the newly reported suppressor CD8+ T cells [37, 10]. The quorum threshold was postulated to be necessary for preventing autoimmunity due to circulating autoreactive T cells without preventing effective T cell responses to pathogens [37]. There is significant empirical evidence that some T cell mediated autoimmune diseases such as MS and T1D are triggered by persistent viral infections. The focus of this paper was to study how persistent or severe infections might cause the breakdown of the mechanism by which collective effects embodied in the quorum model suppresses autoimmunity.

We studied a computational model for T cell development in the thymus, and the response of the resulting T cell repertoire to infection. Upon infection, pathogen-derived peptides are targeted by T cells, and an effective functional response results because the quorum threshold is exceeded to some of these pathogen-derived peptides because they were not encountered by the T cell repertoire during development in the thymus. Our results show that some of the activated T cells that normally exhibit weak cross-reactivity to host-derived pMHC molecules can now be efficiently reactivated by these host antigens in the same tissue because previously activated T cells have a lower activation threshold. Evidence exists for such cross-reactivity between host pMHC molecules and pathogen-derived ones in the case of MS. CD4^+^ T cells have been isolated that are cross-reactive to both EBV-derived peptides and self-antigens like myelin basic protein (MBP), anoctamin 2, alpha-crystallin B, and glial cell adhesion [66, 103, 88, 104, 105, 106]. For MBP, a self-antigen present in myelin sheaths, there is evidence that MS patients have autoreactive T cells that can be activated by APCs presenting EBV peptides [103]. As further evidence, not only is there an increase in the presence of MBP-specific CD8^+^ T cells in MS patients [107], but EBV-specific CD8^+^ T cells isolated from these patients were cytotoxic against cells that were pulsed with MBP and cells transfected to endogenously express MBP [108]. But, our results show that such “molecular mimicry” leads to autoimmunity with low probability for typical infections when just a few pathogen-derived peptides are targeted by the T cell repertoire. For autoimmunity to develop, the T cells activated by the pathogen-derived peptides that are crossreactive to a self-antigen must be sufficiently large in number such that, when added to other T cells that are activated by the same self-antigen, the quorum threshold is exceeded. When just a few pathogen-derived pMHC molecules are targeted, the probability of this happening is small. In other words, peripheral tolerance mechanisms that determine the quorum threshold are able to suppress autoimmunity upon typical infections. This concept is related to the observation that CD8+ suppressor T cells expand upon infection to kill activated autoreactive T cells [8] and shows that, if the balance is tilted toward Tregs, they expand and effector cells do not [10].

Our results also describe how this mechanism goes awry upon persistent or severe infection. In this case, larger numbers of pathogen-derived pMHC molecules are targeted by the T cell repertoire [64, 65]. The T cells that target each of these pMHC molecules has a chance of being weakly cross-reactive to a host pMHC molecule expressed in the same tissue. So, as the number of targeted pathogen-derived pMHC molecules increases, the total probability of activating T cells cross-reactive to the pathogen-derived pMHC and a self-antigen grows. Our results show that this, in turn, results in increasing the chance that the quorum threshold is exceeded by autoreactive T cells and T cells activated by pathogen-derived pMHC molecules that are weakly cross-reactive to a self-antigen. Thus, we find that the probability of autoimmunity being triggered grows with the number of pathogen-derived pMHC molecules targeted during persistent infection [Fig. 6].

Our model brings together the concepts of molecular mimicry and collective effects to provide a consistent explanation for why viral infections such as EBV do not always result in autoimmunity, why certain viral infections trigger certain autoimmune diseases, why there is a lag time between establishment of a persistent viral infection and onset of the corresponding autoimmune condition and why this lag time exhibits large variations. Only certain viruses will result in APCs displaying immunogenic pMHC molecules that activate T cells that are crossreactive to particular self-antigens. This is why certain viral infections are more likely to induce certain autoimmune conditions (e.g., EBV and MS). Our results show that these crossreactive T cells will contribute to exceeding the quorum threshold for a self-antigen to result in autoimmunity only with a finite probability; therefore, a persistent viral infection will only lead to a corresponding autoimmune disease only with some probability [Fig. 6]. One of our findings is the importance of rare peptides that can activate many T cells. If a pathogen-derived peptide triggers many T cells, the chance that T cells cross-reactive to a self-antigen are activated and contribute to exceeding the quorum threshold with respect to self-antigens also increases. Such events are rare [Figs. 6 and Supp. 5] and so autoimmunity may not occur for a very long time, if ever. However, the chance that such an event occurs increases with time during a persistent infection as the chance of T cells targeting more pathogen-derived peptides increases, thus providing an explanation for the lag time. As the lag time is dependent on a stochastic event occurring, it exhibits large variations across individuals.

Our results also show that collective effects embodied in the requirement that the quorum threshold must be exceeded for a functional response lowers the probability of triggering autoimmunity because of variations in thymic selection across individuals, and when infection leads to strong perturbations [Fig. 7]. For example, we found that collective effects embodied in the quorum model make the immune system more robust to autoimmunity when the number of self-antigens T cells are exposed to in the thymus is lower. This situation can be realized either stochastically or due to certain HLA alleles, such as one associated with T1D binds self-peptides less stably. The increase in the probability of triggering autoimmunity upon T cells encountering fewer host-derived peptides in the thymus is reduced by the existence of collective effects and the quorum threshold [Fig. 7A]. However, the influence of collective effects in suppressing autoimmunity does not matter much beyond a threshold value of the quorum threshold, Q [Fig. 7]. The quorum number, Q, is set by the number (or density) of Tregs and other suppressor cells. Thus, this result from our model suggests that beyond a point having more Tregs will not add to robustness of the immune system against triggering autoimmunity.

Inflammation and T cell-mediated cell killing leads to tissue damage, which promotes presentation of self-epitopes that would otherwise not be presented. It has been postulated in the case of T1D that, through this mechanism, also known as epitope spreading, self-reactive T cells are recruited and can initiate an autoimmune response [109]. For example, persistent EV infection leads to inflammation and destruction of islet cells by T cells [110] and an increase in inflammatory cytokines (IFN) [111]. Increase in IFN leads to HLA class 1 over-expression in islet cells and cell death. These factors contribute to increased vulnerability to CD8^+^ T cell bystander activation and targeting of novel self-antigens [112]. More generally, inflammation leads to various effects, such as enhanced crosspriming, etc, that can result in the presentation of self-antigens that are typically “hidden”. This concept of epitope spreading contributing to autoimmunity has also been noted in the context of MS. EBV has been suggested to induce an antiviral immune response against infected cells in the CNS which leads to the release of sequestered self-antigens [113]. In addition, it has also been suggested that EBV-infected autoreactive B cells mediate an autoimmune attack by increasing self-antigen presentation and activating autoreactive CD4^+^ T helper cells that can activate these B cells [114]. In the context of our model, the increased presentation of self-peptides is reflected as an increase in N_T_. Our results show that increasing N_T_ increases the number of T cells activated by self-antigens [Supp. 4], thus contributing to exceeding the quorum threshold. However, consistent with the role of collective effects making the immune system more robust against onset autoimmunity, our results show that the effect of epitope spreading on triggering autoimmunity is mitigated by the requirement that the quorum threshold be exceeded for a functional immune or autoimmune response [Fig. 7B]. Note also that inflammation and more cell death has a higher chance of resulting in expression of rare self-pMHC molecules that can activate many T cells [Fig. 8], thus exceeding the quorum threshold.

The results we have described provide a unifying framework to understand various phenomena, but the model requires further testing. We make a specific prediction to test the veracity of our model. We carried out calculations to identify peptides derived from the proteome of *mycobacterium tuberculosis* that are homologs of peptides from the proteome of C57BL/6 (B6) mice; both sets of peptides bind to the I-Ab MHC molecule expressed by these mice. We predict that immunizing these mice with increasing numbers of the identified *mycobacterium tuberculosis* peptides should result in a higher chance of inducing autoimmune disease. We also expect that the homologous *mycobacterium tuberculosis* peptides that have more hydrophobic amino acids at the sites that contact the TCR should serve as proxies for rare peptides that trigger more T cells, and so we also identify these peptides. Immunizing with a larger number of homolog *mycobacterium tuberculosis* peptides that include these more hydrophobic ones is predicted to increase the probability of triggering autoimmunity. We hope that these experiments will soon be done as they will likely shed new light on the aberrant regulation of peripheral tolerance mechanisms.

We emphasize again that the purpose of our model is not quantitative recapitulation of a specific experimental result. Rather, our goal was to develop a model that could explore the significant consequences of collective effects that are important for a functional T cell response on mitigating the danger of autoimmunity, and how these tolerance mechanisms can be mis-regulated upon persistent or severe infections. Our model has several limitations. In this manuscript, we have focused on the initial stages of T cell activation and proliferation. It is recognized that the affinity of self-reactive T cell responses also determines whether activated clones upregulate tissue homing integrins and induce MHC-I cross-priming of self-antigens [115, 22]. As details of these processes are further quantified, second-generation computational models will incorporate these attributes to better understand when overt autoimmunity ensues, or a given proliferative burst simply expands self-reactive clones. We do not explicitly treat the effects of innate immunity and inflammation, and implicitly treat their effects on increasing self-antigen presentation by increasing the parameter in our model (N_T_) that represents the number of self-antigens presented in the infected tissue. We do not treat Tregs and suppressor CD8+ T cells explicitly either. Their effects are embodied in the quorum threshold number, Q. As we have noted earlier, the value of Q is not going to be a constant as we have assumed, but rather it will be larger for self-peptides that bind more avidly to T cells as interactions with these self-peptides in the thymus are more likely to result in the development of Tregs [93]. Accounting for Tregs and suppressor CD8+ T cells and concomitant differences in the values of Q for different peptides is an important next step for developing a better understanding of the dynamic instability that results in breaking the quorum threshold and initiating autoimmunity. This will require the development of a dynamical version of our model of infection, which is an important next step.

## Github

Link to github with source code and tuberculosis peptide files: https://www.github.com/ pinkzephyr/autoimmune_model

## Acknowledgements

This research was supported by the Ragon Institute of MGH, MIT, and Harvard. RY was also supported by a National Science Foundation Graduate Research Fellowship. MK acknowledges support from NSF grant DMR-2218849. SM was supported by a Physics of Living Systems Fellowship at MIT. We are very grateful for fruitful discussions with Marc Jenkins, Shiv Pillai, and Ruslan Medhzitov.

## Supplementary Materials

### Supp. 1

Amino acid frequencies used to generate TCRs:

| Amino Acid | Homo Sapiens |
| --- | --- |
| C | 0.0225 |
| M | 0.0215 |
| F | 0.0359 |
| I | 0.0434 |
| L | 0.0985 |
| V | 0.0598 |
| W | 0.0123 |
| Y | 0.0263 |
| A | 0.0692 |
| G | 0.0658 |
| T | 0.0536 |
| S | 0.0836 |
| N | 0.0360 |
| Q | 0.0481 |
| D | 0.0476 |
| E | 0.0718 |
| H | 0.0261 |
| R | 0.0568 |
| K | 0.0576 |
| P | 0.0636 |

### Supp. 2

#### E_n_ gap

Dependence of the E_n_-E_p_ gap on E_n_ for the 4% probability of emerging from thymic selection:

**Figure 9:**
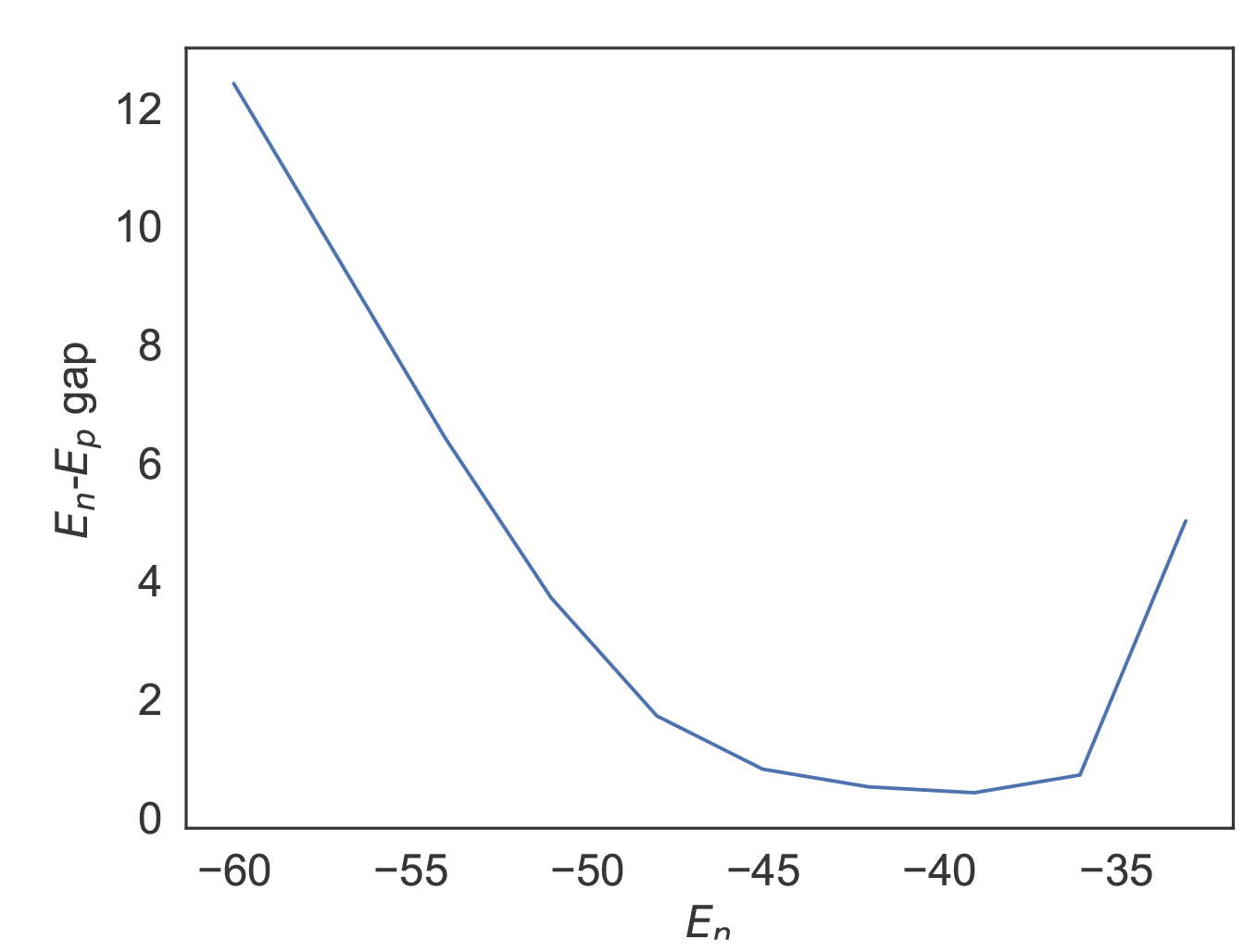
Graph showing the relationship between the value of E_n_ and the corresponding gap between the positive and negative selection thresholds such that there is 4% survival of immature thymocytes during selection.

#### E_n_ dependency

**Figure 10:**
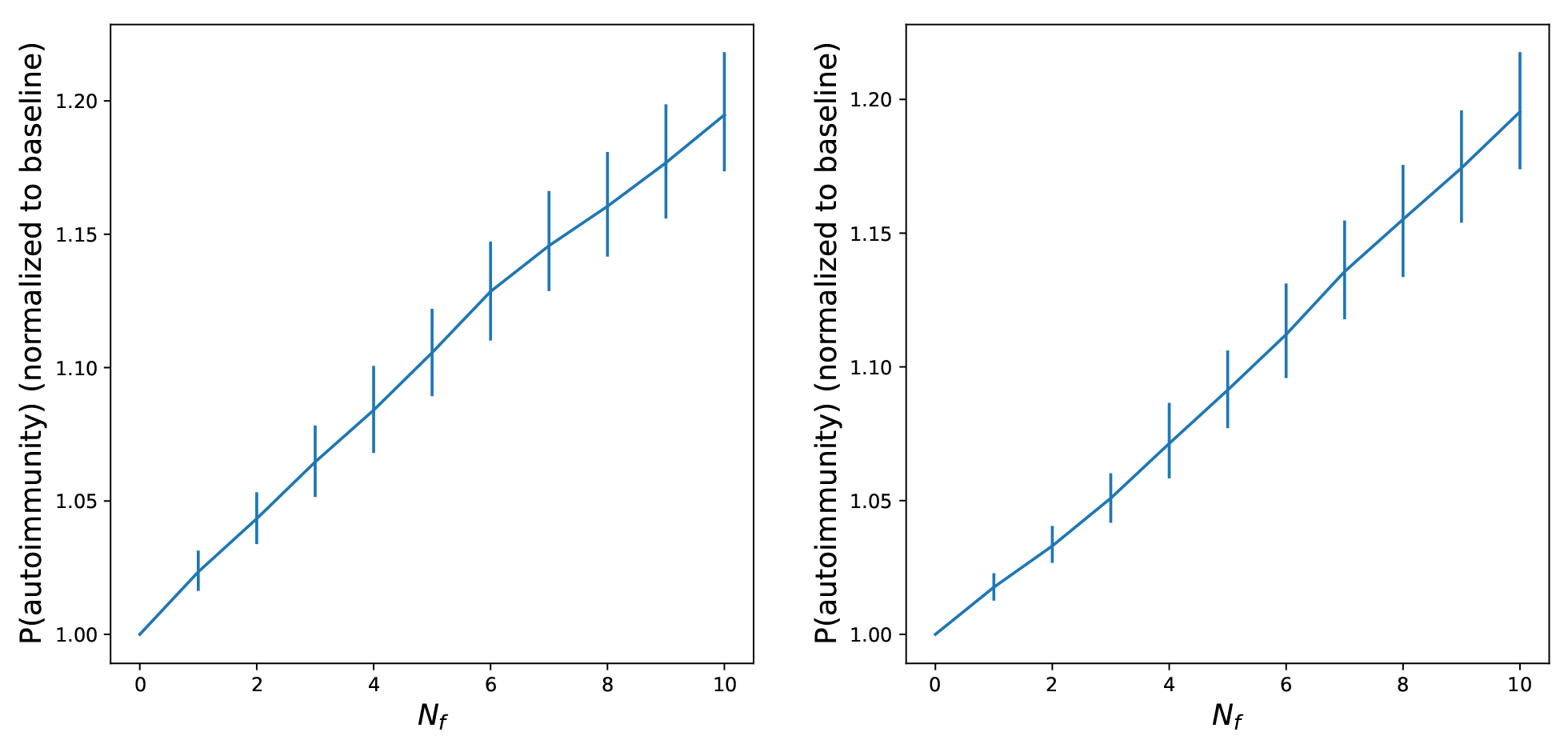
Results of simulations for two additional values of E_n_, -37k_B_T and -39 k_B_T . The results are qualitatively the same as in Figure 6.

### Supp. 3

Graph of activation by P5R peptide:

**Figure 11:**
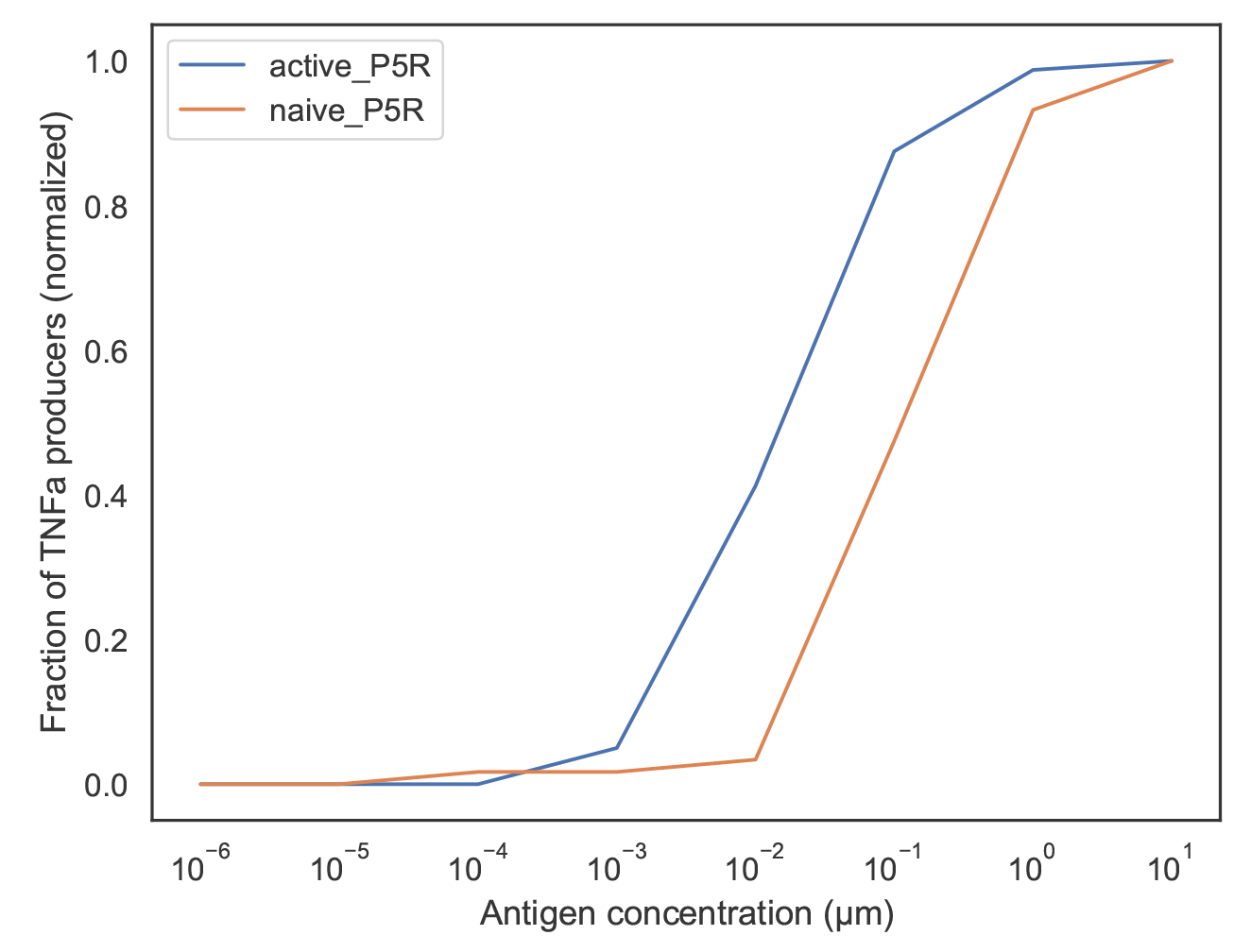
Graph of antigen concentration versus % of T cell population that are TNFα producers when exposed to another antigen, P5R. The blue and red curves are the responses of activated and naive T cells.

### Supp. 4

**Figure 12:**
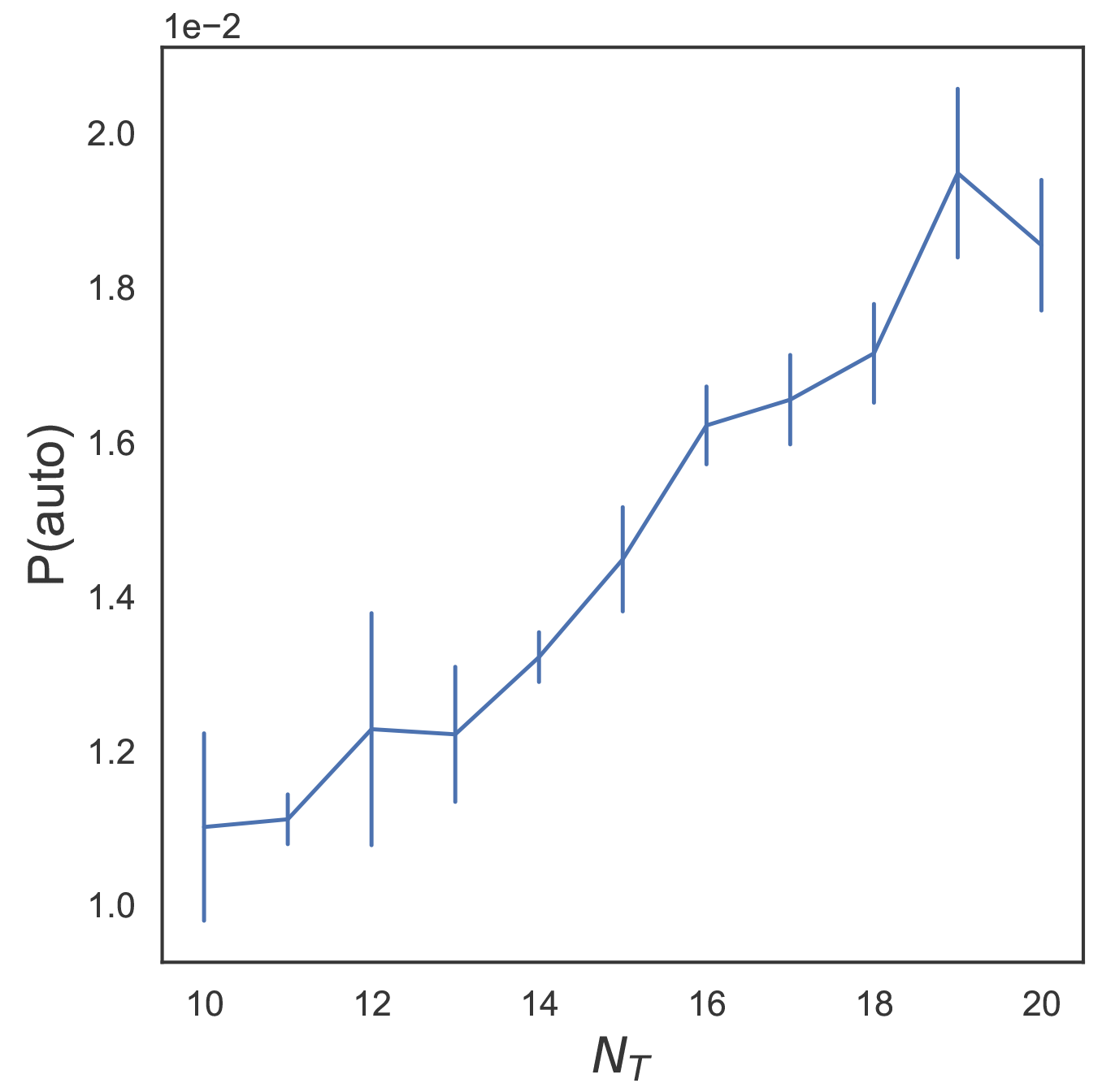
Graph of probability of autoimmunity for increasing N_T_. Simulation parameters are the same as in Table 1, with P(autoimmunity) drawn from N_f_ = 10. Error bars were calculated based on the number of simulations noted in Figure 4.

### Supp. 5

#### Hydrophobicity and Rare Peptides

Let each thymocyte i be characterized by a reactivity parameter, α_i_ *∼* N(0, 1), and each peptide j be identified by a reactivity parameter β_j_ drawn from the same distribution. This reactivity parameter may be thought of as a proxy for hydrophobicity, such that larger values indicate that the thymocyte or peptide is more likely to bind strongly to its counterpart. We can then write the free energy of an interaction between thymocyte i and peptide j as

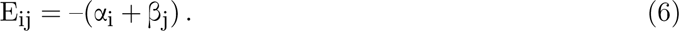

Using this framework, we can simulate selection by generating 1, 000, 000 naive thymocytes, each associated with a reactivity α_i_, and 10, 000 self-peptides associated with a reactivity β_j_. Each thymocyte i encounters a random set, S_j_, of M self-peptides (M = 8,000) during selection, and the mature repertoire consists of thymocytes i where

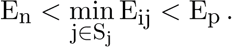

We can then calculate the number of T cells that bind to a sample of 10, 000 self-peptides, and pathogenic peptides, and observe a tail analogous to the one seen in the string model [Fig. 8]. In addition, we can calculate the number of T cells that are activated by pathogenic and self-peptides in the same fashion as the string model, which produces a very similar graph [Fig. 13] as seen by the string model [Fig. 6].

**Figure 13:**
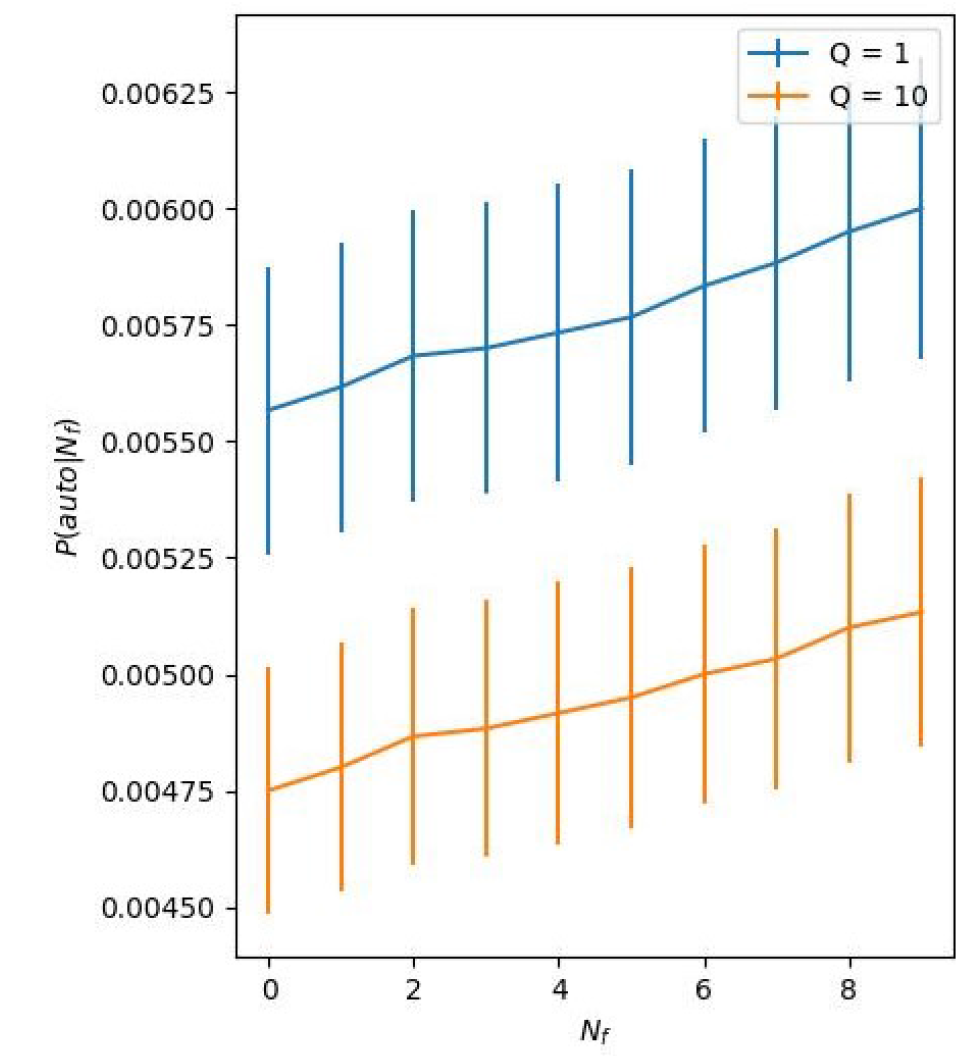
Probability of autoimmunity due to cross-reactivity of pathogen-derived peptides as function of the number of pathogen-derived peptides, simulated based on the coarse grained model described in the text and above. The results are similar to those in Figure 6 in the main text. Parameters used: 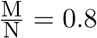, and N_T_ = 10, number of naive T cells T = 10^6^, E_n_ = 2.90, E_p_ = 2.59,

### Supp. 6

Distribution of number of mouse homologs for each tuberculosis peptide:

**Figure 14:**
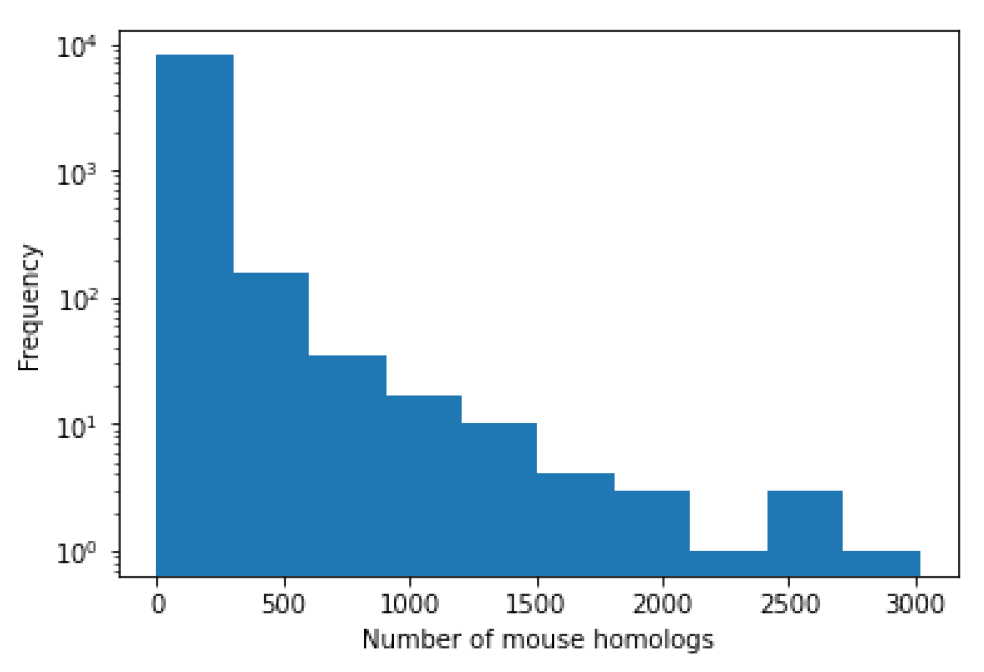
Histogram of mouse homologs per tuberculosis peptide.

**Table S1:**
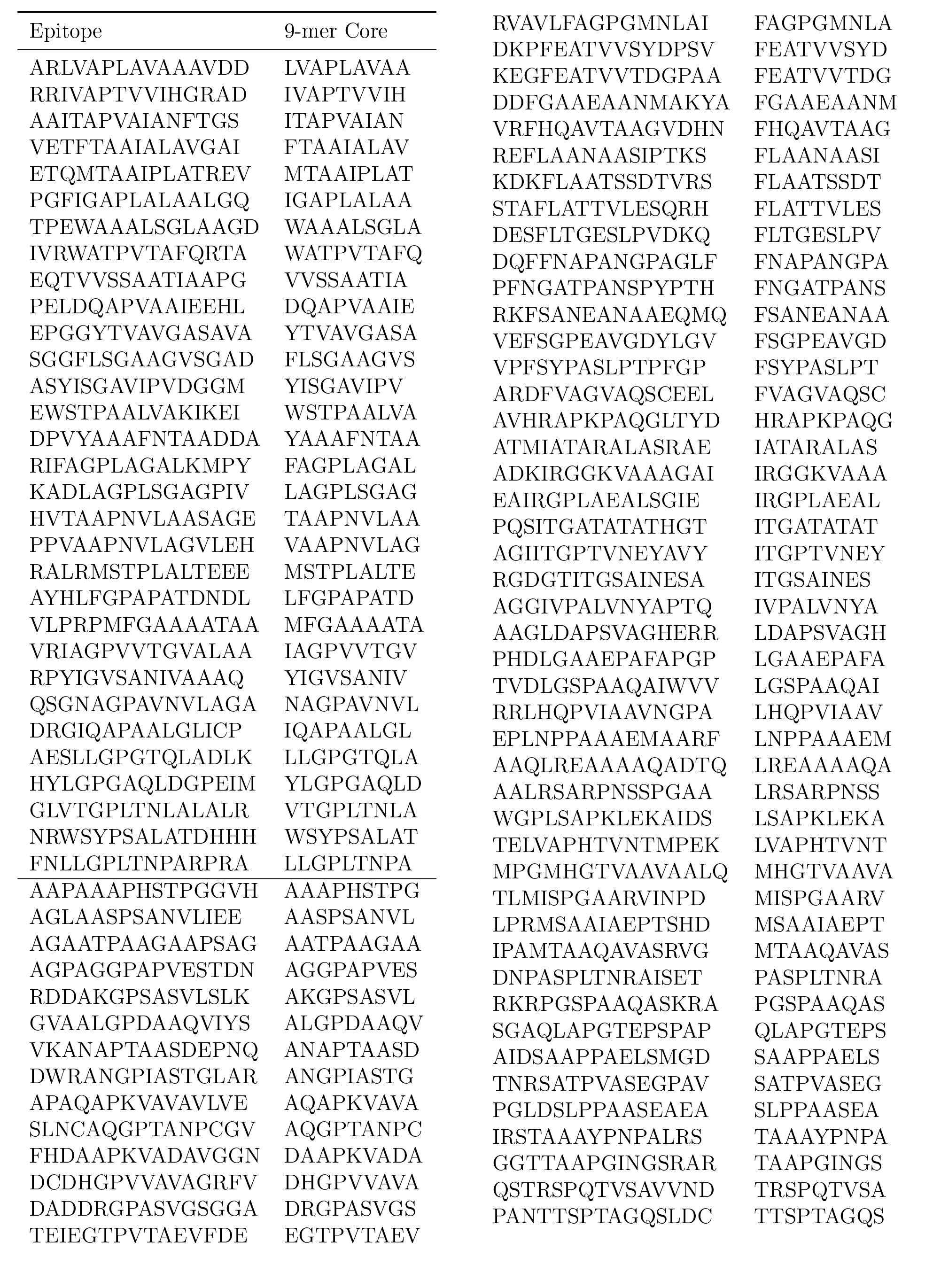
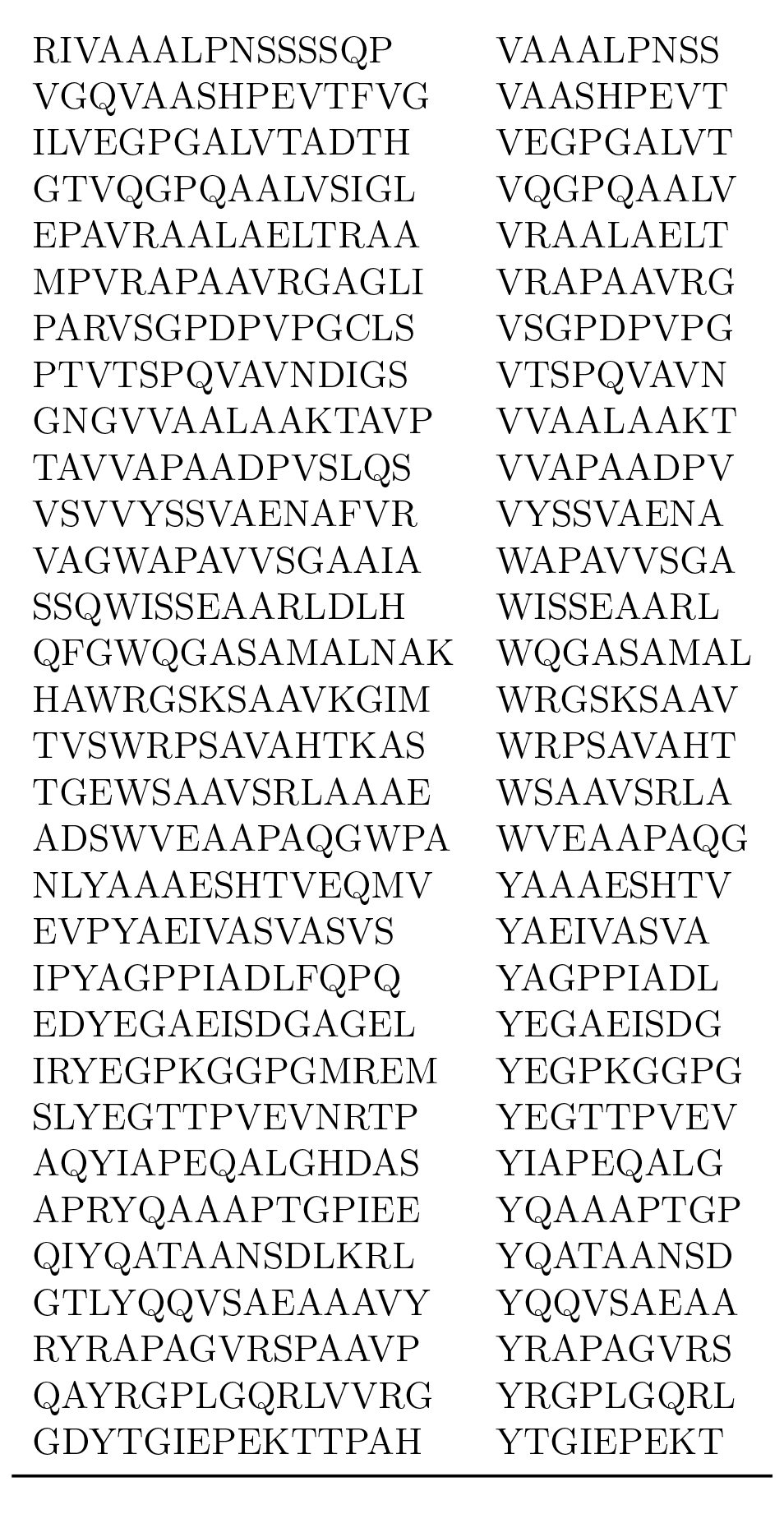
List of tuberculosis epitopes. The first 31 peptides above the line are the most hydrophobic peptides in this list.

